# Integrative plasma lipidomics and proteomics profiling to decipher potential biomarkers of dilated cardiomyopathy

**DOI:** 10.1101/2025.04.13.648582

**Authors:** Shubham Saha, Praveen Singh, Abhi Dutta, Anurag Raj, Mamta Rathore, Deepika Jindal, Hiteshi Vaidya, Santoshi Kumari, Prakash Chand Negi, Shantanu Sengupta, Sandeep Seth, Trayambak Basak

**Affiliations:** School of Biosciences and Bioengineering. IIT-Mandi, HP, India-175075; CSIR-Institute of Genomics and Integrative Biology, New Delhi-110020; Dept. of Cardiology, All India Institute of Medical Sciences, New Delhi-110029; Dept. of Cardiology, Indira Gandhi Medical College & Hospital, Shimla, HP, India-171001

**Keywords:** Dilated Cardiomyopathy, Heart Failure, Biomarker, Lipidomics, Proteomics, scRNA Sequencing, ELISA

## Abstract

**Background:** Dilated cardiomyopathy (DCM), primarily characterised by left ventricular dilatation and systolic dysfunction, is one of the leading causes of heart failure and requires a critical clinical investigative strategy. However, conventional imaging techniques such as echocardiography and MRI, along with some classical CVD markers (NTproBNP, cTnT), fall short in diagnosing DCM-specific phenotypes. Thus, the need for biochemical markers with enhanced accuracy to DCM is of enormous importance. Lipids and proteins play essential roles in maintaining myocardial function. The homeostatic disruption of such biomolecules might contribute to DCM pathogenesis, thus offering to serve as potential biomarkers for DCM. Moreover, the lack of global lipidomics studies and specific protein markers in DCM patients prompted us to explore the disease pathophysiology through an integrative omics-based analysis coupled with an ML-derived approach.

**Objective:** To identify accurate, precise and specific circulatory lipidomic and proteomic biomarkers of DCM using high-resolution mass spectrometry and machine learning (ML)-based approaches.

**Methods:** High-resolution-mass-spectrometry-based lipidomics and proteomics were applied to identify lipid and protein biomarkers in a cohort (n=360) of healthy and DCM patients. Top protein classifiers were further evaluated using single-cell transcriptomics on publicly available datasets from DCM myocardium and validated using ELISA. A biomarker panel was built by the integration of lipidomics and validated proteomics data using machine-learning-based approaches.

**Results:** A total of 125 lipids and 10 proteins have been primarily discovered with significant alterations in DCM (0.8 ≥ FC ≥ 1.2; p_adj_ < 0.05). Using a Boruta-based ML approach, we identified 39 lipids and 10 proteins as primary discriminators between DCM and controls. ELISA validation confirmed the potential of B2M (6.85 ± 2.86 μg/ml vs. 4.26 ± 1.25 μg/ml; p < 0.0001) and Tetranectin (CLEC3B) (1.99 ± 0.88 μg/ml vs. 2.49 ± 0.90 μg/ml; p = 0.0006) to emerge as protein biomarkers of DCM. In line with that, the single-cell transcriptomic analysis showed a similar trend of alteration of tetranectin (CLEC3B) in cardiomyocytes and β2microglobulin (B2M) in varied cell types of the myocardium. Further, Integrated ROC analysis combining the top 8 lipid discriminators with B2M and CLEC3B achieved an AUC of 0.995, demonstrating enhanced diagnostic precision compared to the classical CVD marker NTproBNP (0.965).

**Conclusion:** This study offers a system-omics-based perspective on first-global lipidomic and proteomic changes associated with DCM pathophysiology, with a high potential for diagnostic application.

## Introduction

Dilated cardiomyopathy (DCM) is a type of cardiovascular disease associated with the myocardium, characterised by left or biventricular dilatation and systolic dysfunction ^1^. The dysfunctional myocardium in DCM is caused by the thinning and stretching of the ventricular walls, followed by several pathophysiological changes such as reduced stroke volume and cardiac output, impaired ventricular filling, and an upsurge in end-diastolic pressure leading to progressive heart failure and morbidity^2^. Around 30-50% of total DCM cases are caused by genetic factors and are commonly known as Familial DCM^3^. However, underlying causes of disease onset are still remained unknown in around 60% of cases, classified as idiopathic DCM^4^. The worldwide prevalence of DCM is estimated to be 1 in 250-400 individuals approximately and the incidence is reported to be 5-7 cases per 100,000 individuals annually^5^. As per the Global burden of disease data, there has been a significant surge in DCM incidence in India recently, with cases increasing from 106,460 cases in 1990 to 207,168 new cases by 2019^6^. This sharp increase is concerning and underscores the necessity for proactive diagnostic and preventive strategies^7,8^.

To date, echocardiography or MRI-based imaging techniques have been considered the gold standard method in the diagnosis of DCM^9^. In addition, certain biomarkers such as CRP, BNP, cTnT, MMPs, TIMPs, etc, representing inflammatory, myocyte injury and fibrotic events are also associated with DCM. Such molecules are also useful in diagnosing general cardiovascular disease phenomena^10^. However, such traditional methods of diagnostics cannot accurately delineate the DCM-specific characteristics^10^. Thus, elucidation of circulating biomarkers with specificity to DCM pathophysiological characteristics is essential for effective diagnosis and prognosis of disease-specific phenotypic alterations^7,8^. Lipids are essential for cardiovascular function, offering the majority of its energy requirements while also contributing to cell membrane formation and cell signalling^11^. Concurrently, proteins also ensure the structural and functional integrity of the cardiovascular system. Hence, disruption in the protein and lipid homeostasis might compromise the efficiency of the myocardium, triggering the pathological features of DCM. Identifying key circulating DCM-specific protein and lipid biomolecules can be instrumental in diagnosing idiopathic DCM cases. Previous proteomics analysis of plasma from DCM cohorts have revealed that protein signatures show population specificity and lack consistency across different studies. Such phenomena might be attributed to the small sample size of the discovery cohort, interference of preanalytical variables and lack of prospective cohort-based validations^8^. Furthermore, to the best of our knowledge, no study has examined the global lipidome profile of DCM patients to date^7^. In this scenario, the mass spectrometry-driven omics-based approach has emerged as an efficient tool to characterise biomolecules to elucidate disease-specific markers with more specificity and sensitivity. Such an omics-based approach also possesses the power to discriminate DCM from other cardiovascular diseases with possible phenotypic overlaps. In this study, we employed a high-resolution mass-spectrometry-based approach to profile the proteome and lipidome of the participants. We aimed to identify a panel of potential lipid and protein biomarkers within a cross-sectional cohort, including Indian patients, through a system biology-based methodology. This study has yielded the discovery of 8 lipids and 2 proteins, which have shown remarkable efficiency in diagnosing DCM pathophysiological characteristics.

## Methodology

### Sample collection

Our cross-sectional cohort consists of 360 individuals (188 DCM patients and 172 healthy controls). All the individuals were recruited in the study cohort during their visit to the clinicians at All India Institute of Medical Sciences, New Delhi and Indira Gandhi Medical College, Shimla. The study was ethically approved by the Human Research Ethics Committee of Indian Institute of Technology Mandi, CSIR-Institute of Genomics and Integrative Biology, New Delhi, All India Institute of Medical Sciences, New Delhi and Indira Gandhi Medical College, Shimla (Ethical document no. IIT-Mandi/IEC(H)/2020/24^th^ April/P1, CSIR-IGIB/IHEC/2020-21Dt.23.06.2020, IEC-972/03.10.2020, HFW(MC-II)B(12)ETHICS/2020/12482) and conducted in accordance with the Declaration of Helsinki. All participants provided written informed consent before study entry. All the samples were collected during November 2021-August 2024. Idiopathic DCM patients and corresponding controls sample collections were performed simultaneously. All the participants underwent systematic evaluation procedures including clinician examination, lineage tracing, laboratory biochemical tests and echocardiography. The probability of having familial DCM cases in our cohort has been nullified by performing pedigree analysis. DCM phenotypes were determined according to the recommendation of American Heart Association (AHA). DCM patients were recruited as per the following criteria-1. Left ventricular ejection fraction (LVEF) ≤45% 2. Left ventricular end diastolic diameter (LVEDD)≥55mm. Individuals under the age of 18 years and having a history of ischemic heart disease (IHD), coronary artery disease (CAD), and any other chronic diseases like chronic kidney disease (CKD) and chronic obstructive pulmonary disease (COPD) have been excluded from our cohort. The demographic data and blood samples were collected from the participants at the baseline and before taking any medicines associated with CVD. Blood plasma was isolated from**^(n=^**t**^2^**h**^0^**e^4^**^)^** peripheral blood using venipuncture in the EDTA tubes followed by centrifugation @4⁰C for 15 minutes and frozen immediately at -80°C in different aliquots without thawing until downstream analysis^12^.

### Lipidomics analysis

Lipid extraction: Lipid extraction from the plasma samples was performed using a modified Bligh and Dyer method as described earlier^13^. Briefly, ten microlitres of plasma were used for lipid extraction, it was mixed with 490 µl of ice-cold water (in a glass tube with a PTFE-lined screw cap). This was followed by the addition of 2 ml methanol (MeOH) and 1 ml dichloromethane (DCM). The solution was vortex mixed, monophase was ensured at this stage and this solution was incubated a room temperature for 30 minutes. After incubation, 0.5 ml of water and 1 ml of DCM were again added to this solution, vortex mixed, and centrifuged at 1200 rpm for 10 minutes. The lower layer was collected in a fresh glass tube. 2 ml DCM was again added to the remaining solution in the original extraction tube, vortex mixed, and centrifuged. The lower phase was again collected, and mixed with the first extract. This lipid extract was dried in a vacuum centrifuge by operating it in organic mode at room temperature. The lipid was resuspended in 100 µl of ethanol and transferred to polypropylene auto-sampler vials before LC-MS acquisition.

Data acquisition: Lipidomics data was acquired on a QTRAP 6500+ system coupled to ExionLC UHPLC system. The instrument source gas parameter and chromatographic conditions were used as described earlier (PMID: 35625636). Ten microliters of lipid suspended in ethanol were loaded on the XBridge BEH Amide column (3.5 µm, 4.6×150 mm, Waters) and separation was performed using a binary gradient of buffer A (95% acetonitrile with 10 mM ammonium acetate, pH 8.2) and buffer B (50% acetonitrile with 10 mM ammonium acetate, pH 8.2) in a 24-minute-long method.

QTRAP 6500+ system (SCIEX) equipped with a turbo source with electrospray ionization (ESI) was operated in low mass range mode. Source gas 1(GS1) and source gas 2 (GS2) were set at 50 and 60 psi respectively. Curtain gas was set at 35 psi and the source temperature was set at 500℃. Ion spray voltage was set at 5500 V and -4500 V respectively for positive and negative ion scan modes. Data were acquired in a scheduled MRM (multiple reaction monitoring) method with a variable retention time window and relative dwell time weightage containing an MRM transition library of 1230 lipid species including 12 internal standards, 611 species in positive mode (SM, CE, Cer, TAG, DAG, MAG) and 619 in negative mode (Phospholipids and lysophospholipids). Data acquisition was carried out using Analyst 1.6.3 (SCIEX) and relative quantification was performed in MultiQuant 3.0 (SCIEX).

Lipidomics Data processing and analysis: The lipid species with missing values in more than 30% of samples were filtered out. The data from 204 samples with 1134 lipid species having less than 2% missing values were considered for further analysis: a. Imputation was done for less than 2% missing data in the dataset with median value based on lipid species-specific value across the two groups. Log2 transformation was done after imputation. b. Batch effect correction with age and sex adjustment for 1134 lipid species was done to reduce technical variance using the ComBat function from the SVA package (10.1093/biostatistics/kxj037). Differential expression analysis was performed by a two-sided t-test, where lipid species were found to be altered by more than 1.2 folds with an adjusted p-value of 0.05 between the two groups. The Boruta algorithm using random forest was applied for important feature selection to identify possible lipid markers and iteratively remove lipid species that were statistically less relevant than a random probe between the two groups (10.18637/jss.v036.i11). The ‘Boruta’ function from the R library package was applied to 204 samples with 125 lipids as features, where 39 lipids were finally selected as a putative marker. A two-sample Wilcoxon test with p-value adjustment (BH method) was further applied to show the significant relationship between the categories of identified lipid markers.

### Proteomics analysis

#### Sample preparation

Proteomic studies were performed on the blood plasma samples from the dilated cardiomyopathy (DCM) and healthy control (HC) individuals. Ten microlitres of plasma was diluted with 90 microlitres of 1X phosphate buffer saline (PBS) and protein precipitation from these samples was performed by overnight incubation with pre-chilled acetone followed by centrifugation at 15000g for 15 minutes at 4°C. The protein pellets were suspended in 0.1M Tris-HCl with 8 M urea, pH 8.5 and protein quantitation was performed using the Bradford assay. 20 µg of protein from each sample was processed for quantitative proteomics analysis (SWATH-MS). Protein was reduced by adding a 2mM final concentration of dithiothreitol (DTT) and heating at 56℃ for 30 minutes. Samples were then cooled down at room temperature and cysteine alkylation was performed using a 2.2 mM final concentration of iodoacetamide (IAA) and incubation in the dark, at room temperature for 20 minutes. Samples were then diluted with 0.1 M Tris-HCl buffer, pH 8.5, to bring urea concentration in the sample below 1M concentration before protease digestion. These samples were then subjected to trypsin (V5111, Promega) digestion in an enzyme-to-substrate ratio of 1:20 (trypsin: protein) for 16 hours at 37 ℃. The tryptic peptides were cleaned up using Oasis HLB 1 cc Vac cartridges (Waters) using the manufacturer’s protocol, vacuum dried in a vacuum concentrator, and stored at -20℃ till LC-MS/MS analysis.

#### SWATH-MS data acquisition

The tryptic digests from the samples were suspended in 0.1% formic acid and were analyzed in a label-free proteomics approach on a quadrupole-TOF hybrid mass spectrometer (TripleTOF 6600, SCIEX) coupled to an Eksigent NanoLC-425 system using the Sequential Window Acquisition of All Theoretical Mass Spectra (SWATH-MS) method by operating the mass spectrometer in data-independent acquisition mode. Optimized source and gas parameters were used for the mass spectrometer. 4 μg of peptides were loaded on a trap column (ChromXP C18CL 5 µm 120 Å, Eksigent, SCIEX) and online desalting was performed with a flow rate of 10 µl per minute for 10 min. Peptides were separated on a reverse-phase C18 analytical column (ChromXP C18, 3 µm 120 Å, Eksigent, SCIEX) in a 57-minute-long buffer gradient with a flow rate of 5 µl/minute using water with 0.1% formic acid and acetonitrile with 0.1% formic acid as previously described (PMID: 36237606). SWATH-MS method was created with 100 precursor isolation windows, defined based on precursor m/z frequencies in DDA run using the SWATH Variable Window Calculator (SCIEX), with a minimum window of 5 m/z. Accumulation time was set to 250 msec for the MS scan (400–1250 m/z) and 25 msec for the MS/MS scans (100–1500 m/z). Rolling collision energies were applied for each window based on the m/z range of each SWATH and a charge 2+ ion, with a collision energy spread of 5. The total cycle time was 2.8 seconds.

#### Data analysis

The SWATH-MS raw files were analyzed in Spectronaut 15 software from Biognosys using the directDIA workflow. A Homo sapiens protein database from UniProtKB with 20,396 entries (Swiss-Prot) was used and protein identification was performed with 1% FDR. software search settings were set as follows: trypsin as protease, peptide modification conditions were set as-cysteine carbamidomethylation was set as fixed modification and methionine oxidation and protein N-terminal acetylation were set as variable modifications.The directDIA analysis was performed with default settings; cross-run data normalization was performed where the normalization strategy was set to automatic. Quantitative data was exported in the form of ‘Run Pivot Report’ and differential protein analysis was performed in Microsoft Excel.

#### Proteomics Data Pre-processing

The proteomics dataset comprised 204 samples, with negligible missing values (< 0.03%). The proteins expressed in less than 70% of samples were removed, and the remaining proteins were processed further. To facilitate subsequent analysis, the following pre-processing steps were undertaken. Missing Data Imputation: Missing data were imputed using the median value based on condition-specific measurements within the two experimental groups. This imputation strategy aids in maintaining the integrity of the dataset while mitigating the impact of missing values. Subsequently, a Log2 transformation was applied to normalize the data after imputation. Batch Effect Correction with age and sex adjustment: Technical variance from batch effects in the data was addressed using the ComBat function sourced from the SVA package (10.1093/biostatistics/kxj037) version 3.52.0. This approach ensures the harmonization of the dataset across different batches, enhancing the comparability and reliability of subsequent analyses. A model matrix containing biological covariates, i.e., age and sex, including conditions, was added to the function to adjust their effect.

#### Exploring Differential Expression and Protein Markers

To uncover meaningful insights within the proteomics dataset, a comprehensive analysis plan was executed: Differential Expression Analysis: Leveraging a two-sided t-test, we assessed alterations in protein expression between the two experimental groups. Proteins that exhibited a fold change exceeding 1.2 and a significance level of p_adj_ < 0.05 were identified as differentially expressed, indicating their potential relevance in the context of the conditions. Feature Selection Using Boruta Algorithm: The Boruta algorithm, underpinned by random forest methodology, was employed to facilitate the identification of significant protein markers (10.18637/jss.v036.i11). Boruta is a machine learning algorithm that identifies meaningful features from complex datasets. This method allowed us to find important protein markers by gradually excluding proteins without statistical significance compared to random values in control and DCM groups. It helps to distinguish between actual features (proteins) and shadow features (random copies of the original proteins) and identifies any connections that are not simply due to chance. This comparison is made through many random forests runs and careful statistical tests. In the analysis, the ‘Boruta’ function from the R library package was applied to the dataset encompassing 204 samples and utilizing differentially expressed proteins as features. The Boruta analysis involved multiple random forest runs in which permuted copies of the proteins (referred to as shadow variables), representing proteins with identical distributions as the original proteins but no correlation with DCM, were examined. In each Boruta runs, a set of shadow proteins was created, and random forest models were used to measure how important all proteins are in relation to DCM risk. If a protein’s significance was higher than that of the shadow proteins, we labelled it ‘confirmed’; otherwise, it was ‘rejected’. This process was repeated to remove labelled proteins until all proteins were categorized with a certain number of rounds.

#### Validation of Protein Marker Significance

A two-sample Wilcoxon test was administered to ascertain the significance of the identified protein marker. This test was accompanied by p-value adjustment employing the Benjamini-Hochberg method. By applying this rigorous statistical analysis, we aimed to elucidate the notable relationships between the distinct categories of the identified protein markers.

### Single-cell RNA sequencing data processing and analysis

To gain a detailed understanding of the expression gradients, single-cell transcriptome datasets from transplanted or biopsy samples of DCM and healthy patients were downloaded from the NCBI GEO database (http://www.ncbi.nlm.nih.gov/geo/). UMI counts and associated metadata were obtained from publicly available datasets (GSE109816 and GSE121893). We have employed the Seurat package (V. 3.5.3) in R studio to perform the single-cell transcriptome analysis of DCM hearts and compared it against healthy heart transcriptome 1. First, a Seurat Object was built using the CreateSeuratObject () function and by integrating the matrix data file with the metadata. Quality control metrics were calculated, including the percentage of mitochondrial gene expression. Next, low-quality cells with more than 5% of the mitochondrial genome or transcripts read less than 200 or greater than 2500 were filtered out. The gene expression data is normalised to correct for differences in sequencing depth across cells, using the NormalizeData () function. Highly variable genes were further identified using the FindVariableFeatures () function. Principal Component Analysis (PCA) is performed using the RunPCA () function to reduce the dimensionality of the data.

The filtered cells were clustered based on their gene expression profiles using the nearest-neighbour algorithm with the *FindClusters ()* and *FindNeighbors ()* functions, respectively. The results were visualised with Uniform Manifold Approximation and Projection (UMAP) using the RunUMAP () command. FindAllMarkers () function was used to identify cluster-specific markers genes next. Cluster annotations were done based on their canonical gene markers as described by the parent article 2, we visualised the “genes of interest”.

### Validation of 3 Boruta selected proteins through ELISA

Based upon the discovery phase proteomics, among the 10 Boruta selected proteins, three of the top four proteins were subjected to further validation by Enzyme-linked Immunosorbent assay (ELISA). A case-control cohort consisting of 76 DCM patients and 80 healthy individuals was recruited and plasma samples were collected with informed consent from all the participants. For the quantification of 3 Boruta selected proteins, commercially available sandwich and competitive ELISA kits were used (B2M, *E-EL-H2188, Elabsciences*; CLEC3B, *CSB-EL005531HU, Cusabio*; APOL1, *CSB-EL001940HU, Cusabio*). The concentration of the 3 proteins was compared between DCM and healthy individuals and differences were evaluated using Student’s t-test. Additionally, the correlation was assessed between the plasma level of these 3 proteins with the LVEF of the DCM patients in a quartile-based statistical analysis. ROC curve analysis was also performed to determine the diagnostic performance of each protein and AUC values have been reported.

### Logistic regression model and ROC curve analysis

Logistic regression models were developed to assess the predictive capability of proteomic and lipidomic signature variables in classifying DCM. The models included confounding covariates such as age, sex, BMI, hypertension, and diabetes, implemented using the ‘glm()’ function. Separate models were evaluated for lipids-only, proteins-only, and a combined lipid-protein analysis. The predictive performance of these models, as well as individual lipid and protein variables, were assessed using receiver operating characteristic (ROC) curves and quantified using the area under the curve (AUC) metrics. All analyses were conducted using R (v.4.3.2).

## Results

### Clinical Characteristics

The clinical characteristics of the 204 individuals in the studied cohort are displayed in Table 1. Study participants were recruited between October 2021 to July 2023 with idiopathic DCM patients (n=188), and healthy control subjects (n=172). Out of these, 112 idiopathic DCM and 92 healthy controls were utilized for discovery phase proteomics and lipidomics analysis. The average ages were 47.4±13.7 years and 36.3±12.9 years in the case of DCM and controls respectively. Both the groups were similar in weight, height, body mass index and heart rate. However, the systolic and diastolic blood pressure of the DCM patients (115.3±20.5; 77±13) varied significantly compared to the controls (123.1±15; 83±10.4). Two study groups were compared with respect to multiple serum biomarkers. Creatinine, urea, uric acid and total cholesterol levels of DCM patients varied significantly from the controls (1±0.4 vs 0.8±0.2, P<0.0001; 34.1±16.7 vs 20.7±7.2, P<0.0001; 6.9±2.4 vs 5.5±1.4, P<0.0001; 161.8±41.3 vs 175.8±34.8, P<0.05). Serum NT pro-BNP and cardiac troponin T levels were also significantly increased in DCM patients compared to the controls (1668.4±2871.4 vs 35.2±39.4, P<0.0001 and 18.1±39.3 vs 4.6±1.8, P<0.01). There was a significant decrease in LVEF in the DCM group with respect to the controls (29.7±10.9 vs 60.6±4.7, P<0.0001).

For the validation of altered plasma proteins identified from discovery phase analysis, a total of 156 individuals (76 DCM and 80 controls) were evaluated. The DCM patients and healthy individuals in this cohort differed significantly in terms of their age (41.9±12.7 vs 29.7±7.6, P=1.46E-10), fasting blood sugar level (113.5±53.4 vs 89.2±24.8, P=0.001), serum creatinine (1.1±0.6 vs 0.8±0.2, P=2.96E-05), sodium (138.1±4.4 vs 140.2±3.2, P=0.001), urea (39.7±17.8 vs 21.9±6.8, P=2.90E-12), uric acid (7.2±2.4 vs 5.3±1.4, P=8.34E-08), c reactive protein (7.1±10.7 vs 2.4±3.5, P=0.001) and LDL (88.5±34.8 vs 102.4±28.1, P=0.01). DCM patients were found to have a circulatory NTproBNP level of 3598.6±4959.9 pg/ml. DCM patients were also diagnosed with a significant increase in circulatory cTnT level (15.9±19.7 vs 5.4±3.5, P=0.0001) and substantial reduction in LVEF (27.2±11.0 vs 60.9±2.5, P=8.1E-42) compared to the healthy individuals.

|  | Discovery Cohort |  |  | Validation Cohort |  |  |
| --- | --- | --- | --- | --- | --- | --- |
|  | Control<br>(n=92) | DCM<br>(n=112) | p Value | Control<br>(n=80) | DCM<br>(n=76) | p Value |
| **Baseline** |  |  |  |  |  |  |
| Age (years) | 36.3±12.9 | 47.4±13.7 | 1.58E-08 | 29.7±7.6 | 41.9±12.7 | 1.46E-10 |
| Male/Female | 52/36 | 66/47 | 0.92 | 50/30 | 55/21 | 0.19 |
| Height (cm) | 162.2±8.9 | 161.9±9.7 | 0.83 | 167.5±8.2 | 162.6±12.6 | 0.02 |
| Weight (kg) | 68±10.6 | 65.3±13.8 | 0.12 | 66.7±13.4 | 62.1±12.2 | 0.06 |
| Heart rate | 82.9±11.6 | 85.8±15.6 | 0.16 | 77±10.5 | 84.5±14.6 | 0.0005 |
| Systolic Blood Pressure (mmHg) | 123.1±15 | 115.3±20.5 | 0.002 | 117.7±15.2 | 103.2±19.3 | 1.80E-06 |
| Diastolic Blood Pressure (mmHg) | 83±10.4 | 77±13 | 0.0005 | 74.3±12.2 | 70.1±14.3 | 0.06 |
| BMI | 25.9±3.9 | 25.8±12.2 | 0.97 | 23.7±3.9 | 24.1±6.3 | 0.69 |
| NYHA Class |  |  |  |  |  |  |
| I | NA | 37 (33.04%) | NA | NA | 3 (3.9%) | NA |
| II | NA | 50 (44.6%) | NA | NA | 62 (81.6%) | NA |
| III | NA | 6 (5.4%) | NA | NA | 5 (6.6%) | NA |
| **Comorbidity factors** |  |  |  |  |  |  |
| Hypertension | 18 (19.6%) | 22 (19.6%) | 0.99 | NA | 8 (10.5%) | NA |
| Diabetes | 5 (5.4%) | 26 (23.2%) | 0.0004 | NA | 15 (19.7%) | NA |
| **Serum Biomarkers** |  |  |  |  |  |  |
| Fasting blood sugar (mg/dL) | 97.5±41.9 | 108.4±52.3 | 0.14 | 89.2±24.8 | 113.5±53.4 | 0.001 |
| HbA1c (%) | 5.4±2 | 5.7±2.3 | 0.33 | 5.4±1.1 | 5.8±2.2 | 0.16 |
| Creatinine (mg/dL) | 0.8±0.2 | 1±0.4 | 1.06E-06 | 0.8±0.2 | 1.1±0.6 | 2.96E-05 |
| Sodium (mmol/L) | 139.6±2.4 | 139.3±3 | 0.59 | 140.2±3.2 | 138.1±4.4 | 0.001 |
| Potassium (mmol/L) | 4.7±0.4 | 4.8±0.6 | 0.11 | 4.6±0.5 | 4.6±0.7 | 0.57 |
| Urea (mg/dL) | 20.7±7.2 | 34.1±16.7 | 3.48E-10 | 21.9±6.8 | 39.7±17.8 | 2.90E-12 |
| Uric acid (mg/dL) | 5.5±1.4 | 6.9±2.4 | 3.08E-06 | 5.3±1.4 | 7.2±2.4 | 8.34E-08 |
| C reactive Protein (mg/L) | 3.8±9.3 | 8.2±19.3 | 0.07 | 2.4±3.5 | 7.1±10.7 | 0.001 |
| Total Cholesterol (mg/dL) | 175.8±34.8 | 161.8±41.3 | 0.03 | 172.9±31.7 | 164.3±41.2 | 0.16 |
| Triglyceride (mg/dL) | 140.6±73.5 | 152.4±59.6 | 0.29 | 134.7±66.7 | 152.7±65.6 | 0.11 |
| HDL (mg/dL) | 45±13 | 41.3±10 | 0.06 | 43.6±10.2 | 41.3±11.3 | 0.20 |
| LDL (mg/dL) | 98.9±34.1 | 91.9±28.6 | 0.19 | 102.4±28.1 | 88.5±34.8 | 0.01 |
| VLDL (mg/dL) | 28.1±14.7 | 31.6±13.8 | 0.14 | 26.7±13.3 | 30.5±13.1 | 0.09 |
| NT pro BNP (pg/mL) | 35.2±39.4 | 1668.4±2871.4 | 3.13E-07 | NA | 3598.6±4959.9 | NA |
| Troponin T (pg/mL) | 4.6±1.8 | 18.1±39.3 | 0.003 | 5.4±3.5 | 15.9±19.7 | 0.0001 |
| **Echocardiographic parameters** |  |  |  |  |  |  |
| LVEF | 60.6±4.7 | 29.7±10.9 | 3.67E-61 | 60.9±2.5 | 27.2±11.0 | 8.1E-42 |
| LVEDD | NA | 58.7±7.8 | NA | NA | 60.2±11.5 | NA |
| **Medications** |  |  |  |  |  |  |
| ACEI/ARB/ARNI (%) | 11 (11.9%) | 99 (88.4%) | <0.00001 | NA | 73 (96.1%) | NA |
| β-blocker (%) | 6 (6.5%) | 98 (87.5%) | <0.00001 | NA | 68 (89.5%) | NA |
| Diuretics (%) | NA | 73 (65.2%) |  | NA | 70 (92.1%) | NA |
| Statins (%) | 7 (7.6%) | 25 (22.3%) | 0.004 | NA | 11 (14.5%) | NA |

### High-resolution Mass-Spectrometry-based lipidomic screening reveals unique lipidomic signatures in DCM

A total of 1134 lipids were quantified in the study cohort. Among the 1134 lipid species analysed, 125 have shown significant alteration (0.8≥FC≥1.2 with an adjusted p-value<0.05) between DCM patients and healthy individuals. Out of these 125 lipid species, 8 were significantly increased in the plasma of DCM patients, and 117 lipids were significantly decreased compared to the healthy individuals as depicted in the e volcano plot (Figure 2A). The fold changes of the increased lipid species range from 1.3 to 1.5, with PS(18:0/18:1) having the maximum fold change among the increased lipids. Similarly, the fold changes of the decreased lipids were in the range of 0.5 to 0.8, with PS(14:1/14:1) having the maximum decrease in the plasma of DCM patients.

**Figure 1:**
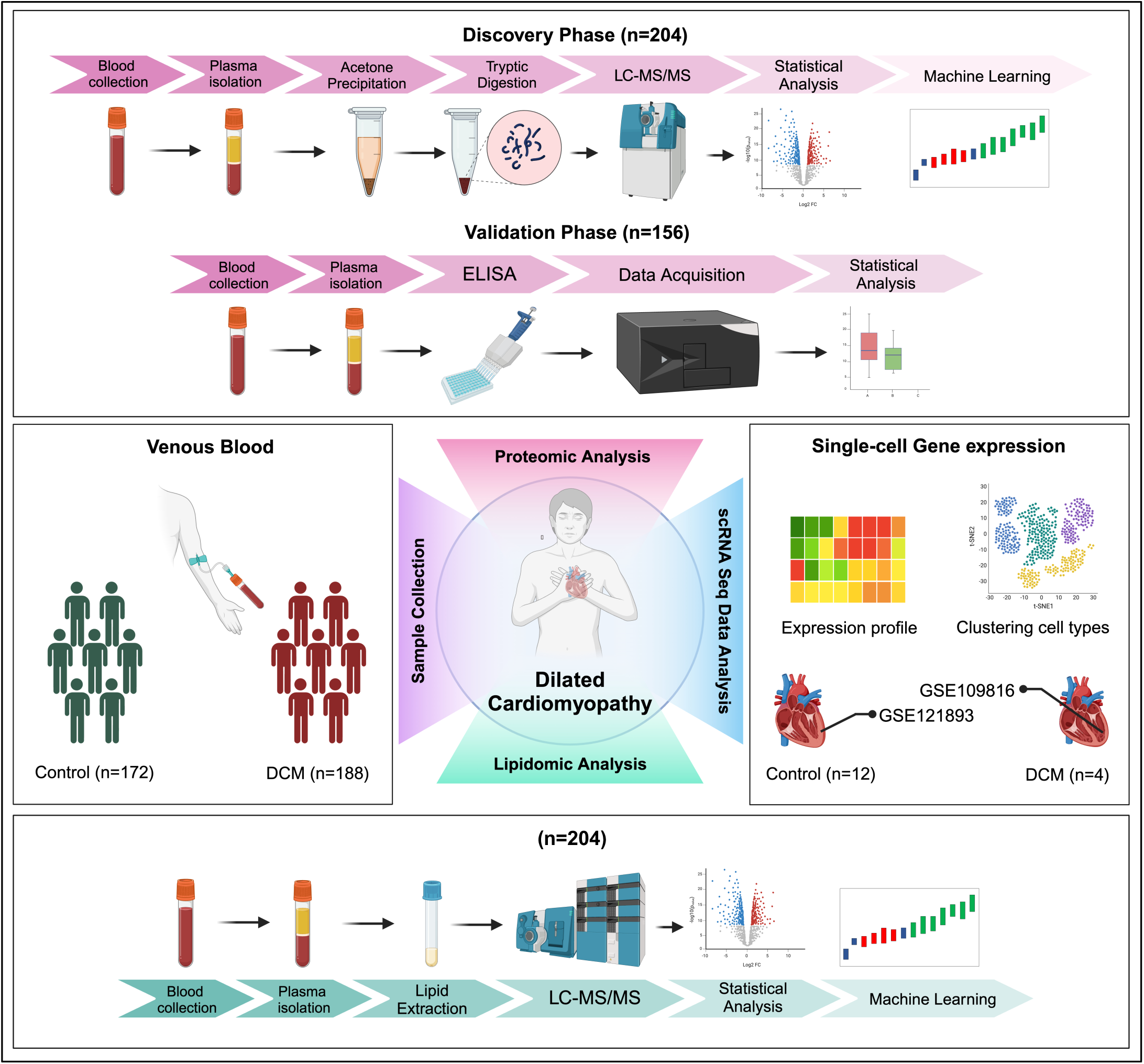
Investigation Framework and Methodology. A total of 360 samples have been analysed through to explore differentially altered lipids and proteins between DCM patients and healthy controls. Some of top proteins have been validated through the ELISA-based method. Gene expression patterns of the top proteins at the single-cell level have also been analysed using available data from cardiac tissue. LC=Liquid Chromatography, MS/MS=Tandem Mass-Spectrometry, ELISA=Enzyme Linked Immunosorbent Assay.

**Figure 2:**
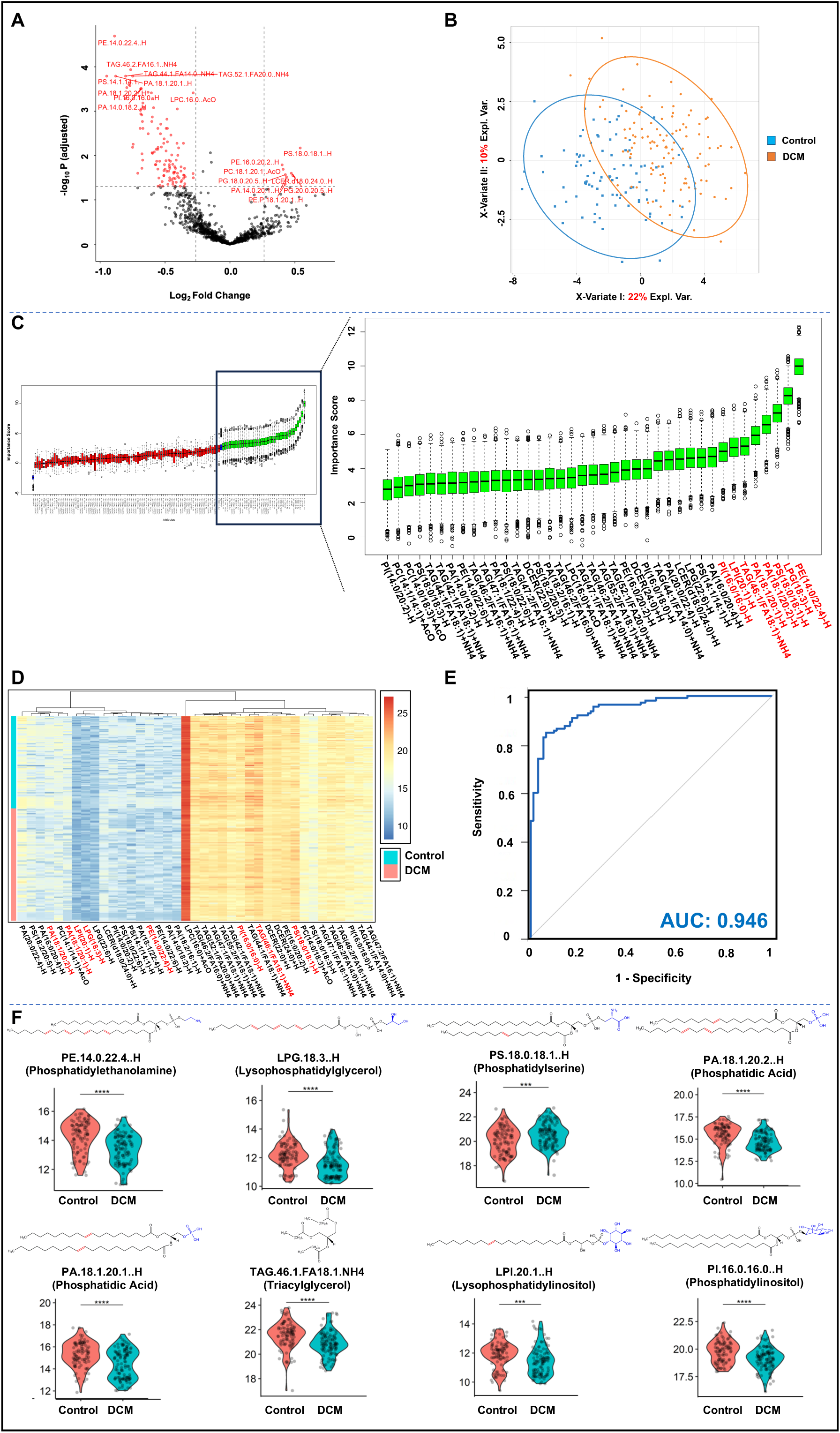
Differentially altered lipids between DCM and healthy controls. A) Volcano plot showing the significantly altered lipids in the DCM patients compared to the controls (0.8≥log2 FC≥1.2, Padj<0.05). B) PLS-DA plot depicting the significant discrimination between DCM and healthy control groups. C) Important lipid feature selection among the significantly altered lipids using Boruta analysis. 39 lipid features selected as important, represented in green. 86 lipids are classified as unimportant, depicted in Red. Shadow features are represented in Blue. D) Heat map representation of Boruta selected lipid features. The colour scale shows the relative intensities of the lipids across DCM and control samples. Red and Blue explains the increase and decrease of the individual lipid species respectively. E) ROC curve analysis predicting the diagnostic accuracy of top 8 Boruta selected lipids individually and in combination. AUC=Area under the Curve. F) Violin plot illustrating significant differential expression of top 8 Boruta selected lipids between DCM and healthy controls along with the chemical structure. ns=Non-significant, *, p<0.05, **, p<0.01, ***, p<0.001, ****, p<0.0001.

### Boruta analysis spotlights critical lipid variables steering DCM pathophysiology

To identify the most relevant lipid species discriminating DCM patients from healthy individuals, the Boruta algorithm was applied to 125 key differentially expressed lipids. This analysis focuses on picking the most important features by trading off the actual variable with their scrambled shadow counterparts. Out of 125 lipids, 39 have been identified as important features. These lipids show consistent relevance in discriminating between the DCM and healthy control groups. Among these 39 lipid species, only 3 are significantly increased and 36 are significantly decreased in plasma of DCM patients (Supplementary Table 1). To further assess the ability of the Boruta-selected lipids to discriminate DCM patients from healthy individuals, partial least squares discriminant analysis (PLS-DA) was applied to maximise the separation between the two groups. The model successfully differentiated the DCM from the control group. The X-variate I has been able to consider 21% of the total variance. An additional 9% variance has been explained by the X-variate II. Together 30% of the explainable variance has been captured by these two components, demonstrating the model efficiency in group separation. However, partial overlap is present between the control and DCM groups (Figure 2B).

Among the important lipid features, PE(14:0/22:4), LPG(18:3), PS(18:0/18:1), PA(18:1/20:2), PA(18:1/20:1), TAG(46:1/FA18:1), LPI(20:1) and PI(16:0/16:0) secure the top 8 positions respectively based on the median importance score cut-off of ≥5 (Figure 2C). These lipids have been predicted to have a role in mitochondrial damage, disruption of calcium homeostasis, induction of ROS production, and activation of necrotic and apoptotic pathways. The differential alteration of 39 Boruta-selected lipids between DCM patients and healthy controls has been illustrated through heatmap (Figure 2D). Individual levels of distribution of these 8 pivotal lipid species in DCM patients compared to healthy controls are shown in Figure 2E. (Supplementary Figure 1). These graphical representations emphasize the potential relevance of the Boruta-selected lipids in the context of DCM pathophysiology. Further, we also assessed, whether statin consumption can contribute to the alteration of these 8 lipid species in DCM patients. The lipid profiles of the DCM patients consuming statin were compared with those of patients who did not consume statin. It has been found that there is no significant difference between the two groups in terms of the lipid profile when the top 8 lipid species were considered (*Supplementary* Figure 3). Hence, it is evident that though statin plays an effective role in altering the LDL cholesterol level, it did not affect the newly discovered DCM-specific lipid biomarkers.

### ROC curve analysis with Boruta selected lipids to elucidate the most relevant lipid variables

Receiver Operating Characteristic curve analysis was performed to evaluate the diagnostic precision of the top 8 lipid biomarkers ((PE(14:0/22:4), AUC=0.827; PA(18:1/20:2), AUC=0.814; PI(16:0/16:0), AUC=0.804; LPG(18:3), AUC=0.795; TAG(46:1/FA18:1), AUC=0.794; LPI(20:1), AUC=0.78; PA(18:1/20:1), AUC=0.786; PS(18:0/18:1), AUC=0.764).). Supplementary Figure 2 represents the ROC curves for the top 8 Boruta-selected lipids. To enhance the diagnostic accuracy, the ROC of the top 8 lipids were combined using a multivariate model to demonstrate the improved performance over individual lipids. This combined model yielded an AUC of 0.946, showing greater sensitivity and specificity (Figure 2E). This highlights the essentiality of utilizing a lipid panel rather than a single marker to improve the diagnostic performance of the lipid markers in DCM.

### Discovery of new circulatory proteomic biomarkers in DCM using Data-Independent-Acquisition based Mass-Spectrometry approach

We have identified a total of 367 circulatory proteins from the discovery cohort using a data-independent-acquisition-based mass-spectrometry approach. Among them, 237 proteins have been identified with 2 or more unique peptides. 50 more proteins have been removed because they could not be identified in at least 70% of the total samples. Moreover, 6 keratins have also been removed to eliminate contaminant interference. Finally, we have proceeded with 181 protein species for further quantitative and statistical analysis. Figure 3A illustrates the volcano plot, which has shown 5 proteins namely β2micoglobulin (B2M), Lipopolysaccharide binding protein (LBP), Haptoglobin (HP), Sushi Domain Containing 1 (SUSD1) and Complement Factor H Related 2 (CFHR2) to be significantly increased in the DCM group compared to the controls along with 5 significantly decreased proteins namely Tetranectin (CLEC3B), Coagulation factor IX (F9), Apolipoprotein L1 (APOL1), Haemoglobin subunit alpha 1 (HBA1) and Haemoglobin subunit beta.

**Figure 3:**
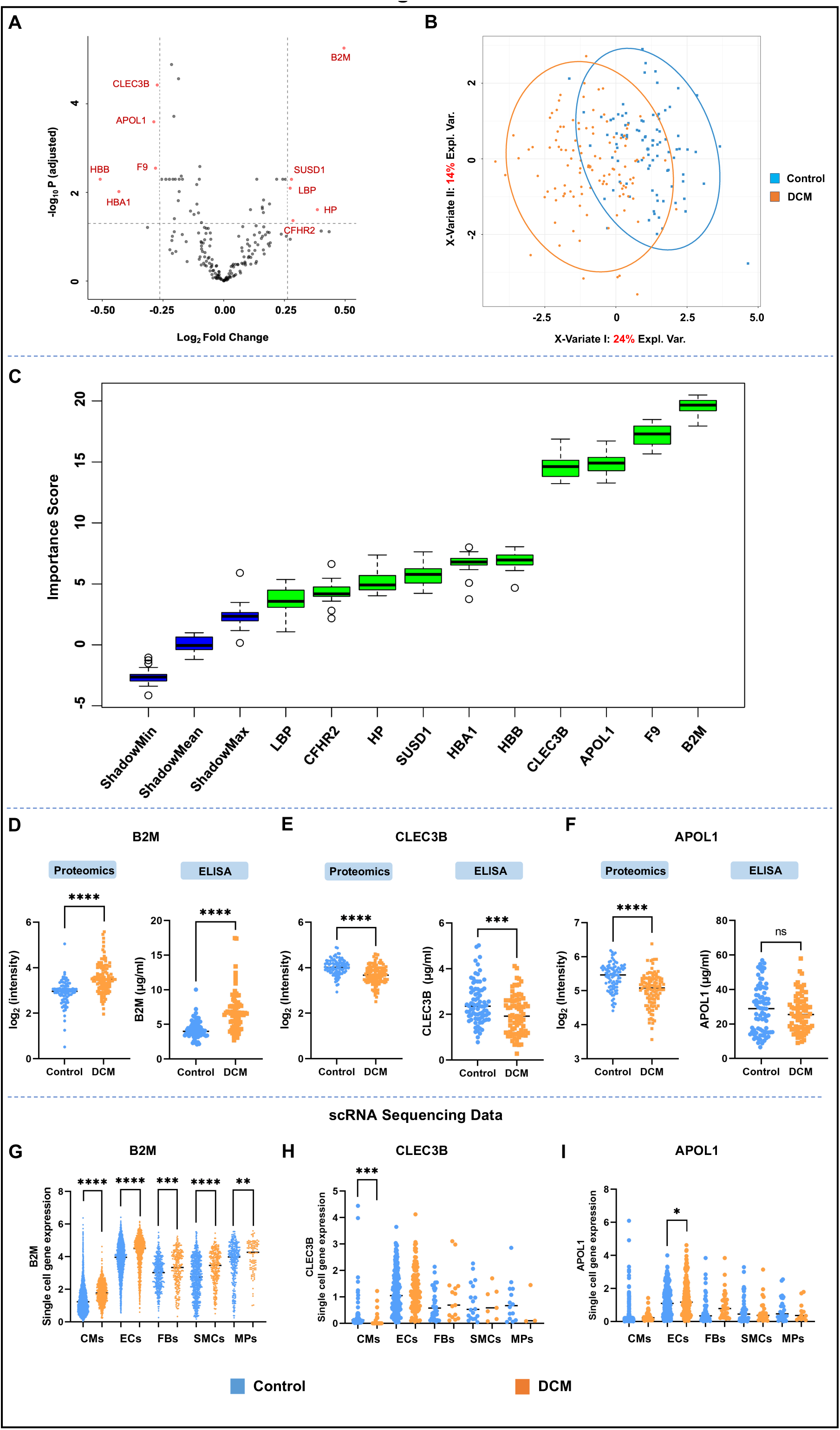
Significantly altered proteins between DCM and healthy controls. A) Volcano plot showing the significant alterations of proteins in the DCM patients compared to the controls (0.8≥log2 FC≥1.2, Padj<0.05). B) PLS-DA plot indicating the classification between DCM and healthy control groups. C) Important discriminatory protein features selection from the significantly altered proteins using Boruta analysis. 10 protein features selected as important, depicted in green. Shadow features are represented in Blue. (D)(E)(F) Comparative analysis between the proteomics and ELISA data of (D) B2M, (E) CLEC3B, and (F) APOL1. (G)(H)(I) Single cell RNA sequencing data analysis of 3 Boruta selected proteins from the cardiac tissue samples explaining gene expression of (G)B2M, (H) CLEC3B and (I) APOL1. CM=Cardiomyocyte, EC=Endothelial Cell, FB=Fibroblast, SMC=Smooth Muscle Cell, MP=Macrophage. ns=Non-significant, *, p<0.05, **, p<0.01, ***, p<0.001, ****, p<0.0001.

### Boruta-driven feature selection identifies essential variables for the discrimination of DCM and healthy controls

Further, to evaluate statistically important differentially expressed proteins between the two groups Boruta algorithm was applied using random forest in the discovery cohort analysis (n=204) revealing 10 significantly (adjusted p<0.05) altered proteins. This analysis yielded all 10 proteins as important features qualified as potential protein biomarkers of DCM (Figure 3C, Supplementary Table 2). Further, Partial Least Squares Discriminant Analysis (PLS-DA) was performed to discern proteomic differences between two groups, DCM and Controls. Figure 3B illustrates the PLS-DA score plot, where each point represents an individual sample. X-variate I and II have accounted for 24% and 14% of variance respectively, a total of 38% of the total explainable variance (Figure 3B). Significant discrimination with partial overlap is observed between the DCM and control groups, indicating a distinct proteomic profile.

### Establishing the proteomic biomarker panel identified from the discovery proteomics analysis

To validate the findings of discovery phase proteomics, we performed an ELISA-based assay with the plasma samples of DCM patients and healthy individuals in another cross-sectional cohort of 156 individuals. A total of 3 proteins (beta-2-microglobulin, Tetranectin and ApoL1) were considered among the top 4 boruta-selected proteins based on a median importance score cut-off of 10. Sandwich and competitive ELISA were performed to quantify the circulatory levels of CLEC3B, B2M and APOL1, respectively, in DCM and healthy controls. As shown in Figure 3D, plasma B2M concentration was significantly higher in the DCM patients compared to the healthy individuals (6.85μg/ml ±2.86μg/ml vs 4.26μg/ml±1.25μg/ml; p<0.0001). In contrast, the plasma CLEC3B level of DCM patients was considerably lower compared to healthy individuals (1.99μg/ml ±.88μg/ml vs 2.49μg/ml±0.90μg/ml; p=0.0006) (Figure 3E). Both B2M and Tetranectin echoed the trend shown by the proteomics analysis. However, no significant change was observed between healthy and DCM samples in the case of APOL1 (26.46μg/ml ±10.51μg/ml vs 28.42μg/ml±13.62μg/ml; p=0.3135) (Figure 3F). Moreover, to decipher the physiological relevance of these key proteins, we have segregated the DCM samples into three quartiles based upon the LVEF (LVEF≤20%, 20%<LVEF≤30% and LVEF>30%) and compared the plasma levels of those proteins in each quartile with the healthy samples as well as other quartiles. Interestingly, the plasma B2M level of the samples in each LVEF quartile was significantly increased compared to the healthy individuals (LVEF≤20% vs Control, 6.78μg/ml ±2.62μg/ml vs 4.26μg/ml±1.25μg/ml, p=0.0001; 20%<LVEF≤30% vs Control, 6.55μg/ml ±2.72μg/ml vs 4.26μg/ml±1.25μg/ml, p<0.0001; LVEF>30% vs Control, 7.55μg/ml ±3.32μg/ml vs 4.26μg/ml±1.25μg/ml, p=0.0003)(Supplementary figure 5). On the contrary, plasma CLEC3B level in each quartile was significantly decreased compared to the healthy samples (LVEF≤20% vs Control, 2.02μg/ml ±0.95μg/ml vs 2.49μg/ml±0.90μg/ml, p=0.0407; 20%<LVEF≤30% vs Control, 1.93μg/ml ±0.89μg/ml vs 2.49μg/ml±0.90μg/ml, p=0.0039; LVEF>30% vs Control, 2.04μg/ml ±0.82μg/ml vs 2.49μg/ml±0.90μg/ml, p=0.0385) (Supplementary figure 5). However, interquartile changes were not significant in any of the cases. Despite remaining unchanged in the validation cohort of DCM and healthy samples, plasma APOL1 concentration was significantly decreased in the LVEF>30% quartile compared to the healthy individuals (22.53μg/ml ±8.55μg/ml vs 28.42μg/ml±13.62μg/ml; p=0.0201). However, plasma APOL1 level was significantly increased in the 20%<LVEF≤30% quartile compared to the LVEF>30% quartile (30.06μg/ml±10.53μg/ml vs 22.53μg/ml ±8.55μg/ml; p=0.0071) (Supplementary figure 5).

Further, we focused on B2M and CLEC3B, demonstrating positive outcomes in the validation cohort and evaluated their correlation with the DCM pathophysiological progress. We have conducted univariate logistic regression analysis. Both B2M and CLEC3B was found to be significantly associated with DCM (for B2M, OR=7.63, CI 95%: 3.99-16.29, P=1.36E10^−8^; For CLEC3B, OR=0.55, CI 95%: 0.38-0.78, P=0.001). Among all the potential confounding factors, age and hypertension have been found to be significantly associated with the DCM with an odds ratio of 4.31 (CI 95%: 2.68-7.53, P= 2.36E-08) and 9.29 (CI 95%: 1.65-174.62, P= 0.038) respectively. However, sex and BMI failed to have a significant correlation with DCM. To account for such confounding factors, we have also conducted a multivariate logistic regression model. After adjustment of age, sex, BMI and hypertension, both B2M and CLEC3B remained an independent predictor of DCM with an adjusted odds ratio of 4.67 (CI 95%: 2.34-10.43, P=0.00005) and 0.51 (CI 95%: 0.31-0.81, P=0.006) respectively. Therefore, plasma levels of B2M and CLEC3B are independently established as two protein biomarkers of DCM.

We have also performed ROC curve analysis for B2M and CLEC3B to establish the robustness of these two potential markers in DCM diagnostics. Both these proteins were found to have a remarkable ability to discriminate between DCM and healthy controls with an AUC value of 0.8636 and 0.8383, respectively. To improve the accuracy of DCM characterisation, both proteins were combined and found to have a cumulative AUC of 0.8896, indicating an enhanced diagnostic performance compared to the individual proteins (Figure 4A, Supplementary Figure 6).

**Figure 4:**
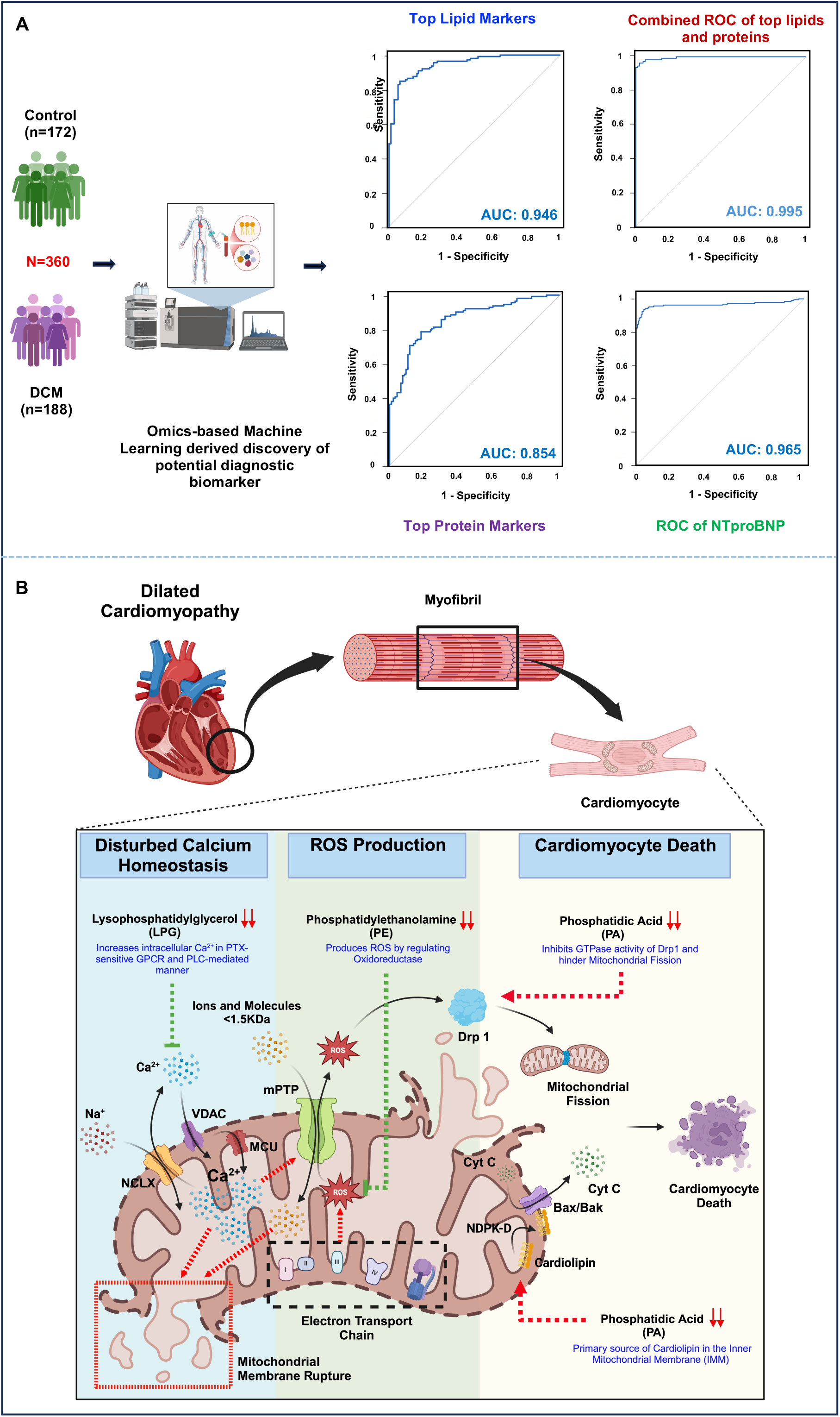
Diagnostic precision of top lipid and protein markers and anticipated mechanistic role of few lipids in DCM pathophysiology. A) Combined ROC curve analysis of top 8 lipids and 2 proteins and comparison with the ROC of NTproBNP. B) Probable association of some of the top lipids with the pathophysiological progress of DCM. Central Illustration: A high-resolution mass-spectrometry-based machine learning-driven approach was adopted to decipher potential lipid and protein biomarkers discriminating the DCM patients from healthy individuals, including a study cohort of 360 individuals. The data acquired from proteomics analysis was correlated with the scRNA seq data of myocardial tissue from a publicly available dataset. Three of the top 4 proteins as per the Boruta Analysis were validated in an independent cross-sectional cohort with an ELISA-based approach. The top 8 lipid species from the Boruta analysis, along with the two validated protein biomarkers, were combined, and an ROC analysis was performed. The combined panel of top lipids and proteins presented an AUC of 0.995, higher than the AUC of classical CVD marker NTproBNP (0.965), indicating better diagnostic precision of the combined panel derived from our study.

### Evaluation of myocardium-specific gene expression of the circulatory proteomic biomaerkers identified in DCM patients

Considering the convoluted aetiology of DCM, we explored the transcriptional profiles of human DCM hearts at the single-cell level to understand the potential alteration of the circulatory biomarkers in the diseased DCM myocardium. Interestingly, B2M has been found to increase significantly in terms of gene expression in the myocardial tissue of DCM patients, and a similar trend has been observed as per the outcome of the plasma proteomic analysis. B2M was also found to be significantly increased in different cell types of the myocardium, such as cardiomyocytes (CM), endothelial cells (EC), fibroblasts (FB), smooth muscle cells (SMC) and macrophages (MP) (Figure 3D). Further, when we tried to dissect the single-cell gene expression profile of Tetranectin (CLEC3B), it was found to be decreased significantly in cardiomyocytes (whereas the rest of the cell types showed no significant changes) of diseased myocardium only reflecting the decreased levels observed in the circulatory levels of DCM patients (Figure 3E). In contrast, APOL1 was found to be significantly increased in the endothelial cells only (Figure 3F). Alongside, we have also explored the single-cell level gene expression patterns of the rest of the Boruta-selected proteins to inspect whether circulatory proteome profiles are consistently reflected within myocardium-specific context. Interestingly, similar to B2M, F9 and HP were also found to show similar trends with the findings of proteomic analysis. The gene expression of F9 was decreased significantly in the myocardial tissue (specifically in cardiomyocytes) of DCM patients. However, HP has shown increased gene expression in the myocardial tissue. Strikingly, all the diseased myocardial cell types have shown significantly higher gene expression in the case of HP (Supplementary Figure 4). Further, it has been observed that alteration in the gene expression of HBA1 was mostly attributed to the endothelial cells, as a significant increase of HBA1 was evident in the single-cell gene expression of the endothelial cells (Supplementary Figure 4). However, HBA1 and HBB gene expression have shown a trend opposing the evidence observed in the proteomic data as both these genes were significantly increased in the myocardial tissue of DCM patients. In line with the evidence furnished by HBA1, endothelial cells of the DCM myocardium have shown a significant decrease in the single-cell level expression of LBP and SUSD1 genes. However, no other cell types have shown such alterations of the gene expression at the single-cell level (Supplementary Figure 4). Notably, CFHR2 has shown no expression in DCM or healthy myocardium.

### Integrated multivariate-logistic regression model using Lipidomics and Proteomics to diagnose DCM

To demonstrate the notable enhancement in diagnostic accuracy, a multivariate logistic regression was built with the top protein (B2M and Tetranectin) and lipid biomarkers (PE(14:0/22:4), PA(18:1/20:2), PI(16:0/16:0), LPG(18:3), TAG(46:1/FA18:1), LPI(20:1), PA(18:1/20:1), PS(18:0/18:1)). This model yielded an enhanced AUC of 0.995 enabling superior sensitivity and specificity for the diagnosis of DCM even compared to classical less-specific NT-pro BNP (AUC=0.96). (Figure 4A). These findings thus emphasise the augmented benefits of combining proteomic and lipidomic datasets to accomplish more credible biomarker panel for improving diagnostic precision.

## Discussion

In this study, we have considered a substantially large cohort to report the first-ever comprehensive analysis of plasma lipidome and proteome in the Indian population. This study helped us explore multiple spectrums: 1) significantly altered lipids and proteins with the potential to discriminate DCM patients from healthy individuals. 2) the most important lipids and proteins among all the altered protein and lipid species using the Boruta algorithm. 3) clustering of samples using PLS-DA analysis along with the predictive value of potential protein markers represented in the form of ROC curve. Moreover, 3 of the top 4 proteins have been validated through ELISA-based methods in order to establish their diagnostic significance. Along with that, 8 top lipids and 2 proteins have been analysed through a combined ROC generation approach, which produced a remarkable AUC of 0.995, exceeding the AUC value of classical CVD marker NTproBNP (0.965). All these analyses thus yielded a panel consisting of 8 lipids and 2 protein species with the potential to screen DCM phenotypes specifically and distinctly from the other forms of cardiovascular diseases sharing phenotypical overlaps with DCM.

Lipids are one of the crucial elements of the cardiovascular system as it serves as the primary source of energy, required for the heart’s functionality^14^. ∼70% of resources that provide energy to the heart consist of fatty acid derivatives. Moreover, lipid modulates many signalling pathways at the molecular level and serves as the primal element for cell membrane formation^14^. Hence, alteration of lipid profile plays a pivotal role in cardiovascular disease pathophysiology. In this study, we explored such lipids affecting cardiovascular health in dilated cardiomyopathy patients through a mass-spectrometry-based analysis of the plasma samples. These lipid features carry the utmost potential to differentiate DCM phenotypes from healthy individuals. Boruta Analysis has shown a significant decrease in lysophosphatidylglycerol (LPG 18:3) species to be one of the top discriminators of DCM and healthy individuals. Lysophosphatidylglycerol is one of the major species among the hydrolyzed phospholipids and comes under the umbrella class of Lysoglycerophospholipids^15^. This class of lipids mostly exist in abundance in plasma or interstitial fluid but is scarce in intracellular environment due to the hyperactivity of lysoacyltransferases. This enzyme enables the incorporation of acyl chain in lysophospholipids and subsequent conversion to corresponding phospholipids^16,17^. However, LPG concentration is very low (0.4μM in Humans; 600-800nM in Mice) in circulation compared to the other lysoglycerophospholipids such as lysophosphatidylcholine (LPC), Lysophosphatidylinositol (LPI), and Lysophosphatidylethanolamine (LPE)^18^. Despite being a low abundant molecule, LPG contributes significantly to the cardio-vascular disease biology. Significant elevation of plasma LPG level in acute coronary syndrome (ACS) patients has been evidenced to establish an association of circulatory LPG with the occurrence of CVD ^19^. There are previous evidence that LPG dictates the Ca^2+^ homeostasis in OVCAR-3 ovarian cancer cells ^20^. LPG stimulates intracellular Ca^2+^ increase through a GPCR-PLC-ERK-AKT signalling cascade in OVCAR-3 cell line^20^. Intracellular Ca^2+^ increase is known to be coupled with varied cellular responses such as apoptosis, one of the crucial phenomena presented in DCM pathophysiological characteristics. However, the question remains whether such kind of molecular caricature will be mimicked in the cardiovascular environment or not, these pieces of evidence indicate a possible interrelationship between molecular signalling driven by LPG species and the pathophysiological progress of DCM (Figure 4B).

Boruta analysis has also yielded multiple circulatory phospholipid species as top discriminators between DCM and healthy individuals. Phospholipids, being the prime structural element of the cell membrane, were also found to contribute significantly to cardiovascular disease progression^21^. In cardiomyocytes, the sarcolemma is majorly constituted by phosphatidylcholine (PC)(45%) and phosphatidylethanolamine (PE)(37%) along with trace amounts of phosphatidylinositol (PI), phosphatidylserine (PS), phosphatidylglycerol (PG) and sphingomyelin (SM)^22^. Our study outcome has confirmed 3 PE lipid species among the top discriminators between DCM and healthy individuals. PE, the second most abundant phospholipid of the sarcolemma, predominantly resides in the inner leaflet of the membranes^23^. Earlier findings indicate a vital role of PE in the differentiation of P19 teratocarcinoma cells into cardiac myocytes as PE elevation has proven to be an early-stage event in the process^24^. Serum lipidomic assessment has revealed PE(12:1e/22:0) to be significantly decreased in post-myocardial infarction heart failure patients with a high predictive value^25^. However, PE(12:1e/22:0) has been found to be negatively correlated with BNP^25^. Another mass spectrometry-based plasma lipidomic study has shown PE to increase in patients with atrial fibrillation^26^. Furthermore, PE supplementation has been found to escalate atrial fibrosis in AngII-induced fibrosis in mice^26^. This study has further explored the inducing role of PE in aggravation of ferroptosis, mitochondrial damage and cell death^26^ (Figure 4B). However, apoptotic events in cardiomyocytes are not only dependent on such factors but also rely upon the PS composition of the cardiomyocyte membrane ^27^. PS generally resides in the inner leaflet of the lipid bilayer and translocation of PS to the outer leaflet is a hallmark of apoptotic or necrotic events in cells^27^. Such relocation of PS triggers the accumulation of pro-apoptotic factors in the mitochondrial membrane and subsequent release of cytochrome c from the intermembrane space to the cytoplasm, thus potentiating the intrinsic pathway of apoptosis^27^. In our study, multiple PS lipid species have been altered in the plasma of DCM patients, reflecting the involvement of apoptotic factors in the pathophysiological progress of the disease. Among all the differentiating PS lipids, circulating PS(18:0/18:1) level has increased and PS(14:1/14:1), PS(18:2/20:5), PS(18:0/22:6) and PS(18:0/18:3) have decreased significantly qualifying among the major differentiators. Such findings demonstrate a substantial correlation between cardiomyocyte membrane lipid composition and cardiac mishaps during the disease progression of DCM demanding the need of deciphering the exact role of these lipids in disease pathophysiology.

Another minor constituent of sarcolemma, namely PI has been found to be associated with ferroptotic affairs in OVCAR-8 (human high-grade serous ovarian carcinoma) and 786-O (human clear cell renal cell carcinoma) cells^28^. Evidence shows that the incorporation of arachidonic acid (AA) into PI increases the susceptibility of cells towards ferroptosis-associated molecular pathways. Such molecular events happen with the intervention of some molecular players such as MMD, ACSL4 and MBOAT7. These proteins contribute to the integration of AA into PI^28^. In our study, we have identified 3 decreased PI species (PI(14:0/20:2), PI(16:0/16:0) and PI(16:0/18:0)) among the key differentiators of DCM . In concordance with our study outcomes, previous evidence also furnished plasma PI levels to be decreased in coronary heart disease patients when compared to the healthy individuals^29^. These findings, as a whole, emphasise the probable involvement of PI lipid species in the onset of pathophysiological characteristics of DCM.

Membrane phospholipids also undergo a chain of metabolic processes to produce an array of by-products which might have some role in CVD pathogenesis. Besides contributing to the membrane integrity, PC lipid species also produce a variety of metabolic derivatives such as phosphatidic acid (PA) which is brought forth by the action of phospholipase D^30^. Our study demonstrates the alteration of some PA lipid species in the plasma of DCM patients as well as establishes their prominence in discriminating DCM cases from healthy individuals. The features include PA(18:1/20:1), PA(18:1/20:2), PA(16:0/20:4), PA(20:0/22:4), PA(18:2/16:1), PA(18:1/22:4) and PA(14:0/18:2) with significant decrease in DCM plasma samples. Though the association of these lipids with DCM pathophysiology is still to be explored, prior research has unveiled the contribution of PA lipids in other forms of cardiovascular diseases such as atherosclerosis and cardiac hypertrophy^31,32^. PA has been found to induce hyperactivity of Na^+^-Ca^2+^ exchanger and subsequent release of Ca2+ from the sarcoplasmic reticulum into the cardiomyocyte cytoplasm^33^. PA also activates PLC (Phospholipase C) to induce cleavage of PIP2 and consequent upregulation of cellular IP3 concentration in fibroblast cells^32^. PA has also been found to facilitate the activation of MAPK and downstream signal transduction to induce hypertrophy in cardiomyocytes^34^. Further findings imply a cyclic interrelationship between PA and cardiolipin^35^. Cardiolipin is produced from PA in the inner mitochondrial membrane (IMM) and partly relocated to the outer mitochondrial membrane (OMM). A small fraction of Cardiolipin translocated to the OMM is further converted to PA by the action of MitoPLD^35^. Additional evidence indicates that mitofusin-mediated mitochondrial fusion machinery is dependent on the hydrolysis of cardiolipin by mitoPLD and subsequent production of PA in the OMM^36^. In parallel, PA makes the mitochondria resistant to fission by interacting with Drp1 and inhibits the GTPase activity of Drp1, so that Drp1 aggregates on the mitochondrial membrane without effective mitochondrial division^37^ (Figure 4B). Whether the alterations of circulatory PA lipid species of our study reflect disarray in the cardiac mitochondrial membrane remains a question for further investigation. However, such complementary evidence suggests a possible association of circulatory PA decrease with aggravated mitochondrial fission and subsequent cardiomyocyte death, a hallmark of DCM pathophysiological progress.

Alongside lipids, proteins are equally critical contributors in the pathophysiological onset of DCM as proteins play a crucial role in the preservation of structural and functional integrity of the cardiovascular system. Proteins dictates the stability of cardiac myocardium by maintaining the structural and functional integrity and executing signal transduction in the cardiovascular cellular-niche. Previously we have reported that none of the population-specific DCM plasma proteomics could identify pathology-specific proteomic biomarkers as these studies were limited with either state-of-the-art mass-spectrometry or lack validation^8^. In our study, we have identified 10 significantly altered plasma proteins through data-independent-acquisition (DIA) based proteomics in conjunction with Boruta-based machine learning models as major distinctive features between DCM and healthy individuals. β2micoglobulin was established as the top discriminator with a significant increase in the plasma of DCM patients which corroborates with its gene expression pattern in the single-cell level. Cardiomyocytes, endothelial cells, fibroblasts, smooth muscle cells and macrophages have been found to express B2M. Significant upregulation was observed in each cell type of DCM heart compared to the healthy ones. ELISA data has also validated the trend of plasma B2M increase in the DCM patients. In fact, B2M was also found to be increased in each of the EF quartiles compared to the controls. However, interquartile changes were not significant. Prior analysis has shown increasing circulating β2micoglobulin concentration to be attributed to Major Adverse Cardiovascular Events (MACE) as estimated by a quartile-based analysis including patients with carotid atherosclerosis^38^. Quartiles with a higher concentration of circulatory β2micoglobulin have also shown greater susceptibility towards myocardial infarction, stroke and death compared to the quartiles with lower blood concentration of β2micoglobulin^38^. Higher concentration of β2micoglobulin in the circulation has been predicted as a potential marker for risk stratification in acute heart failure patients as well^39^. In another prospective cohort-based study, circulatory β2micoglobulin concentration has been estimated as a predictive marker for mortality caused by coronary heart disease (CHD)^40^. Such pieces of evidence imply considerable role of β2micoglobulin in CVD diagnosis and prognosis. Furthermore, a significant increase in β2micoglobulin levels has been observed in the plasma samples of idiopathic DCM patients. In addition, plasma β2micoglobulin concentration has been in a significant positive correlation with the serum T-cell marker and sodium concentration as well as blood urea nitrogen and plasma renin activity. Such study outcomes highlight a possible contribution of exaggerated T-cell response and hyper-stimulated renin-angiotensin-aldosterone activity^41^.

Further, myocardial fibrosis is one of the distinguishing features of the pathophysiological progress of DCM^42^. In cardiomyopathies, replacement fibrosis is mostly visible in the remodelled myocardium^43^. Hyperactivation of inflammatory responses, neurohormonal anomalies and mechanical stress due to apoptosis or necrosis of cardiomyocytes ultimately results in excessive extracellular matrix deposition in the myocardium^44^. This leads to adverse phenotypical expression in cardiomyopathy patients. Though the mechanistic understanding of the onset of fibrosis in cardiomyopathy cases is a subject of further exploration, our study has revealed an association of circulatory tetranectin (CLEC3B) with the disease progression in DCM. Analysis of plasma proteome revealed tetranectin to be significantly decreased in the circulation of DCM patients compared to healthy individuals. This outcome remains in line with previous evidence, where tetranectin was also found to be significantly lower in the serum of HF patients^45^. ELISA analysis has also shown CLEC3B to be decreased in the plasma of DCM patients in a separate cross-sectional cohort. Tetranectin has been shown to have a higher predictive value in comparison with serum BNP in heart failure ^45^. However, serum tetranectin level was found to be negatively correlated with various fibrotic markers such as PICP and PINP ^45^. Interestingly, cardiac tissue tetranectin gene expression assessment produces a contradiction with the trend observed in plasma or serum samples. Tissue tetranectin was found to be significantly higher in HF patients ^45^. Even, tissue tetranectin gene expression was significantly and positively correlated with the gene expression of various fibrotic markers such as collagen I, collagen III, MMP2, MMP9, TIMP1, and galectin-3 ^45^. Such divergence in the tissue gene expression pattern and circulatory profile of tetranectin is possibly due to the compensatory action of the protein in countering interstitial fibrosis following accumulation in the myocardium. Evidence has been furnished in favor of the protective role where exogenous tetranectin was found to protect the neurons from neurotoxicity by inhibiting apoptosis and autophagy, suggesting a potential therapeutic role in neurodegenerative disorders^46^. Moreover, in DCM, the gene expression of tetranectin among cardiomyocytes was found to be significantly downregulated. Tetranectin has been found to protect cardiomyocytes from hypoxia-induced injury by suppressing apoptosis through activation of PI3K/Akt pathway^47^. Nonetheless, the speculative cardioprotective role of tetranectin requires scientific validation through further exploration.

### Study limitations

Our study possesses several positive aspects in terms of the recruitment of study samples with critical characterization of DCM in AIIMS, New Delhi and IGMC, Shimla. Moreover, we have applied an unbiased systems biology-based approach with the inclusion of the first-ever global lipidomics workflow along with a robust and well-established proteomic analysis platform integrated with machine learning-based framework. Further, we have correlated the findings of the proteomics analysis with the outcomes of a publicly available scRNA seq dataset acquired from myocardial tissue samples, which provides us with significant insight into the mechanistic understanding of the potential protein biomarkers. Additionally, we have validated three of the top 4 Boruta-selected protein markers in an independent cross-sectional cohort through an ELISA-based approach, which bolsters confidence in our novel observations along with comprehending their significance in clinical application. Besides the identification of potential lipid and protein biomarkers, their clinical utility has also been assessed in terms of predictive power, diagnostic sensitivity and specificity while taking confounding factors such as age, BMI, hypertension and diabetes into account.

Nonetheless, a few constraints must be considered regarding our study. The study findings might be influenced by population bias since samples were collected from a specific demographic or geographic region. Hence, the study outcome might not apply in a broad spectrum to all DCM populations. Moreover, a prospective cohort-based study would have produced more fruitful outcomes since it allows tracking participants over a longer period and capturing the disease progression, which makes it more robust for studying long-term health outcomes. Further, only a few proteins have been validated, leaving many candidates without experimental confirmation. Besides, functional insights of identified lipids and proteins were mainly inferred from literature-based evidence rather than direct experimental validation, which might lead to over-interpretation or misinterpretation. Hence, besides providing valuable perception regarding DCM pathophysiology using a machine-learning driven multi-omics approach, its limitations highlight the need for further mechanistic validation of lipid and protein markers through cellular and animal models to translate outcomes into clinical applications.

## Conclusion

In summary, plasma lipidomic and proteomic profiling of DCM patients has put forward a novel panel of lipid and protein biomarkers with high predictive value (AUC=0.995) that can be correlated with the distinct phenotypical characteristics associated with DCM. These protein and lipid molecules thus qualify the criteria of potential biomarkers for DCM and hold the capability of therapeutic utility. Further, some of these proteins and lipids may contribute to the pathophysiological progression of the disease by inducing oxidative stress leading to necrosis and apoptosis of cardiomyocytes, as well as functioning as cardio-protective agents by reducing the fibrotic phenomena appearing in the myocardial environment during DCM. However, most of the findings are novel and the contribution of these molecules towards DCM pathophysiology remains unexplored. Thus, this study lay the foundation for future mechanistic investigation to assess the therapeutic potential of these novel protein and lipid molecules in DCM.

## Perspective

Beyond offering potential biomarkers for improved diagnosis of DCM, our study holds remarkable translational potential in terms of DCM risk stratification and further therapeutic targeting. Since the contribution of these lipid and protein biomarkers in DCM disease pathophysiology is poorly understood, our study paves the way for further exploration at the cellular and molecular level.

*Perspective 1:* To increase the translational potential of the lipid and protein biomarkers, a prospective cohort can be integrated to validate the clinical applicability of the biomolecular signature in a larger independent cohort.

*Perspective 2:* The insights gained from the previous literature regarding the molecular functionality of the lipid and protein signature indicate their potential role in DCM pathophysiological progression. Such pieces of evidence suggest the need for validation of these biochemical markers through experimental models (in-vitro and in-vivo) and lay the foundation for exploring precise therapeutic strategies for DCM through mechanistic investigation.

## Acknowledgements

The author sincerely acknowledges all the investigators from IIT Mandi, CSIR-IGIB, New Delhi, AIIMS New Delhi and IGMC, Shimla for their contributions to this study.

## Central Illustration

A high-resolution mass-spectrometry-based machine learning-driven approach was adopted to decipher potential lipid and protein biomarkers discriminating the DCM patients from healthy individuals, including a study cohort of 360 individuals. The data acquired from proteomics analysis was correlated with the scRNA seq data of myocardial tissue from a publicly available dataset. Three of the top 4 proteins as per the Boruta Analysis were validated in an independent cross-sectional cohort with an ELISA-based approach. The top 8 lipid species from the Boruta analysis, along with the two validated protein biomarkers, were combined, and an ROC analysis was performed. The combined panel of top lipids and proteins presented an AUC of 0.995, higher than the AUC of classical CVD marker NTproBNP (0.965), indicating better diagnostic precision of the combined panel derived from our study.

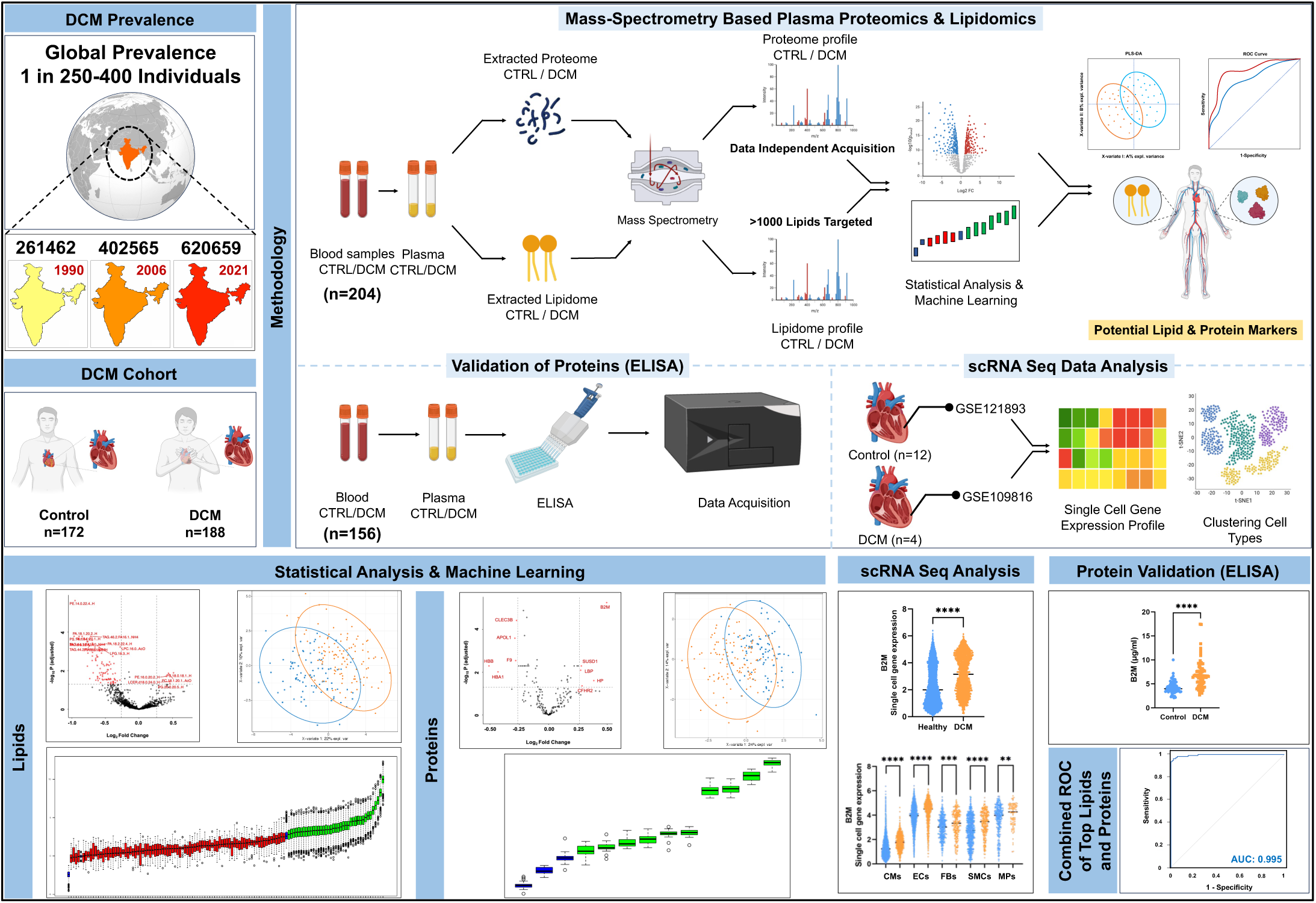

