## Supplementary Information for "Integrative plasma lipidomics and proteomics profiling to decipher potential biomarkers of dilated cardiomyopathy"

### Appendix


| Sl. No | Content | Page No |
|---|---|---|
| 1 | Supplementary Table 1: Boruta-selected differentially altered lipids in DCM patients | 3 |
| 2 | Supplementary Figure 1: Violin plots of Boruta selected lipids showing significant alteration between DCM and Healthy individuals | 4-6 |
| 3 | Supplementary Figure 2: ROC curve analysis of top 5 Boruta selected lipids depicting their predictive accuracy in DCM and Healthy controls | 7-8 |
| 4 | Supplementary Figure 3: Alteration of top 5 Boruta selected lipid species in DCM patients based on their status of Statin consumption | 9 |
| 5 | Supplementary Table 2: Boruta-selected significantly altered proteins in DCM patients | 10 |
| 6 | Supplementary Figure 4: Comparative analysis of the plasma levels of Boruta selected proteins and their tissue expression pattern from single-cell RNA sequencing data | 11-12 |
| 7 | Supplementary Figure 5: Dot-plot representation of ELISA data of 5 Boruta selected proteins and comparative analysis of the plasma levels of those proteins across different LVEF quartiles | 13 |
| 8 | Supplementary Figure 6: ROC curve analysis of 2 validated proteins (B2M and CLEC3B) through ELISA depicting their predictive accuracy in DCM and Healthy controls | 14 |
| 9 | References | 15-16 |

*Supplementary Table 1: Boruta-selected differentially altered lipids in DCM patients*

| Lipid Name | log fc | -log p | -log $p_{adj}$ | Function |
|---|---|---|---|---|
| PC(14:1/14:1) | -0.692 | 4.793 | 2.948 | Defective PC metabolism produces ventricular arrhythmia and mitochondrial dysfunction[1]. |
| PC(14:0/18:3) | -0.392 | 2.332 | 1.292 | |
| PE(14:0/22:4) | -0.952 | 8.683 | 5.634 | Plays role in atrial fibrosis, ferroptosis, mitochondrial damage and cell death [2]. |
| PE(14:0/22:6) | -0.448 | 2.726 | 1.551 | |
| PE(16:0/20:2) | 0.405 | 2.970 | 1.707 | |
| PI(16:0/16:0) | -0.655 | 5.184 | 3.226 | Induces ferroptosis in cancer cell lines [3]. Enzymes associated with PI biosynthesis and metabolism induces cardiomyocyte apoptosis[4], hypertrophy[5], defective excitation-contraction coupling and disrupts $Ca^{2+}$ homeostasis[6]. |
| PI(16:0/18:0) | -0.585 | 3.215 | 1.880 | |
| PI(14:0/20:2) | -0.696 | 4.141 | 2.524 | |
| PS(18:0/18:1) | 0.515 | 3.236 | 1.886 | Relocation of PS from inner leaflet of membrane to outer leaflet potentiates apoptotic and necrotic events by activating intrinsic pathway of apoptosis [7]. |
| PS(14:1/14:1) | -0.966 | 6.128 | 3.778 | |
| PS(18:2/20:5) | -0.643 | 2.618 | 1.504 | |
| PS(18:0/22:6) | -0.482 | 2.670 | 1.530 | |
| PS(14:0/22:5) | -0.499 | 1.758 | 0.924 | |
| PS(18:0/18:3) | -0.515 | 3.711 | 2.253 | |
| PA(18:1/20:2) | -0.847 | 6.214 | 3.778 | Induces hypertrophy, hyperactivity of $Na^+$-$Ca^{2+}$ exchanger, disrupts $Ca^{2+}$ homeostasis in cardiomyocytes [8]. Also activates mitochondrial fusion machinery and inhibits fission-associated factors [9]. |
| PA(14:0/18:2) | -0.693 | 5.519 | 3.425 | |
| PA(18:1/20:1) | -0.892 | 6.178 | 3.778 | |
| PA(16:0/20:4) | -0.772 | 5.667 | 3.506 | |
| PA(18:2/16:1) | -0.641 | 4.157 | 2.524 | |
| PA(20:0/22:4) | -0.733 | 2.797 | 1.587 | |
| PA(18:1/22:4) | -0.588 | 2.816 | 1.587 | |
| LPC(16:0) | -0.280 | 5.161 | 3.226 | Exacerbate ROS production in cardiovascular environment[10]. |
| LPG(18:3) | -0.594 | 5.119 | 3.217 | Disrupt $Ca^{2+}$ homeostasis [11]. |
| LPG(22:6) | -0.484 | 3.747 | 2.271 | |
| LPI(20:1) | -0.390 | 2.486 | 1.402 | Induces $Ca^{2+}$ increase and affects membrane potential[12]. |
| DCER(24:0) | -0.420 | 3.865 | 2.334 | Involved in cellular stress responses, autophagy and apoptotic pathways[13]. Also emerged as a potential markers for diabetes, cancer and neurodegenerative diseases[14]. |
| DCER(22:0) | -0.373 | 3.927 | 2.369 | |
| LCER(d18:0/24:0) | 0.458 | 2.522 | 1.428 | Promotes endothelial dysfunction[15], oxidative stress[16] and left ventricular hypertrophy[17]. |
| TAG(46:1/FA18:1) | -0.629 | 4.566 | 2.796 | Linked to atherosclerotic event, followed by strokes and heart attacks[18]. |
| TAG(44:1/FA14:0) | -0.762 | 5.652 | 3.506 | |
| TAG(46:2/FA16:1) | -0.719 | 6.216 | 3.778 | |
| TAG(44:1/FA18:1) | -0.682 | 3.237 | 1.886 | |
| TAG(52:1/FA20:0) | -0.698 | 5.845 | 3.574 | |
| TAG(47:1/FA18:1) | -0.644 | 4.308 | 2.639 | |
| TAG(46:2/FA14:0) | -0.646 | 4.211 | 2.560 | |
| TAG(46:2/FA16:0) | -0.554 | 3.928 | 2.369 | |
| TAG(55:2/FA18:1) | -0.445 | 3.755 | 2.271 | |
| TAG(47:2/FA16:1) | -0.625 | 4.398 | 2.711 | |
| TAG(42:1/FA18:1) | -0.543 | 3.058 | 1.780 | |

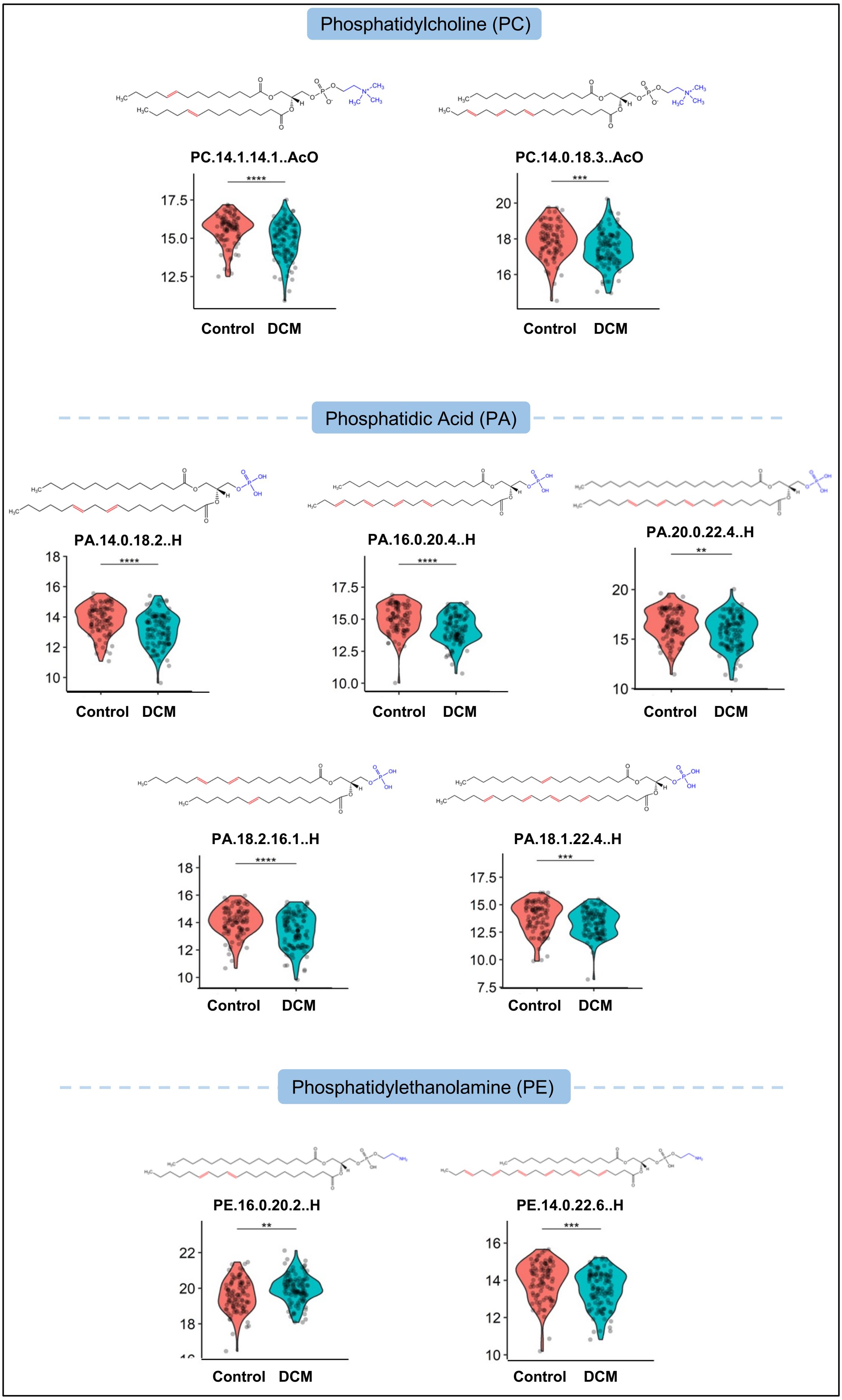

Phosphatidylcholine (PC)
PC.14.1.14.1..AcO
****
17.5
15.0
12.5
Control DCM
PC.14.0.18.3..AcO
***
20
18
16
Control DCM
Phosphatidic Acid (PA)
PA.14.0.18.2..H
****
18
16
14
12
10
Control DCM
PA.16.0.20.4..H
****
17.5
15.0
12.5
10.0
Control DCM
PA.20.0.22.4..H
**
20
15
10
Control DCM
PA.18.2.16.1..H
****
18
16
14
12
10
Control DCM
PA.18.1.22.4..H
***
17.5
15.0
12.5
10.0
7.5
Control DCM
Phosphatidylethanolamine (PE)
PE.16.0.20.2..H
**
22
20
18
Control DCM
PE.14.0.22.6..H
***
16
14
12
10
Control DCM

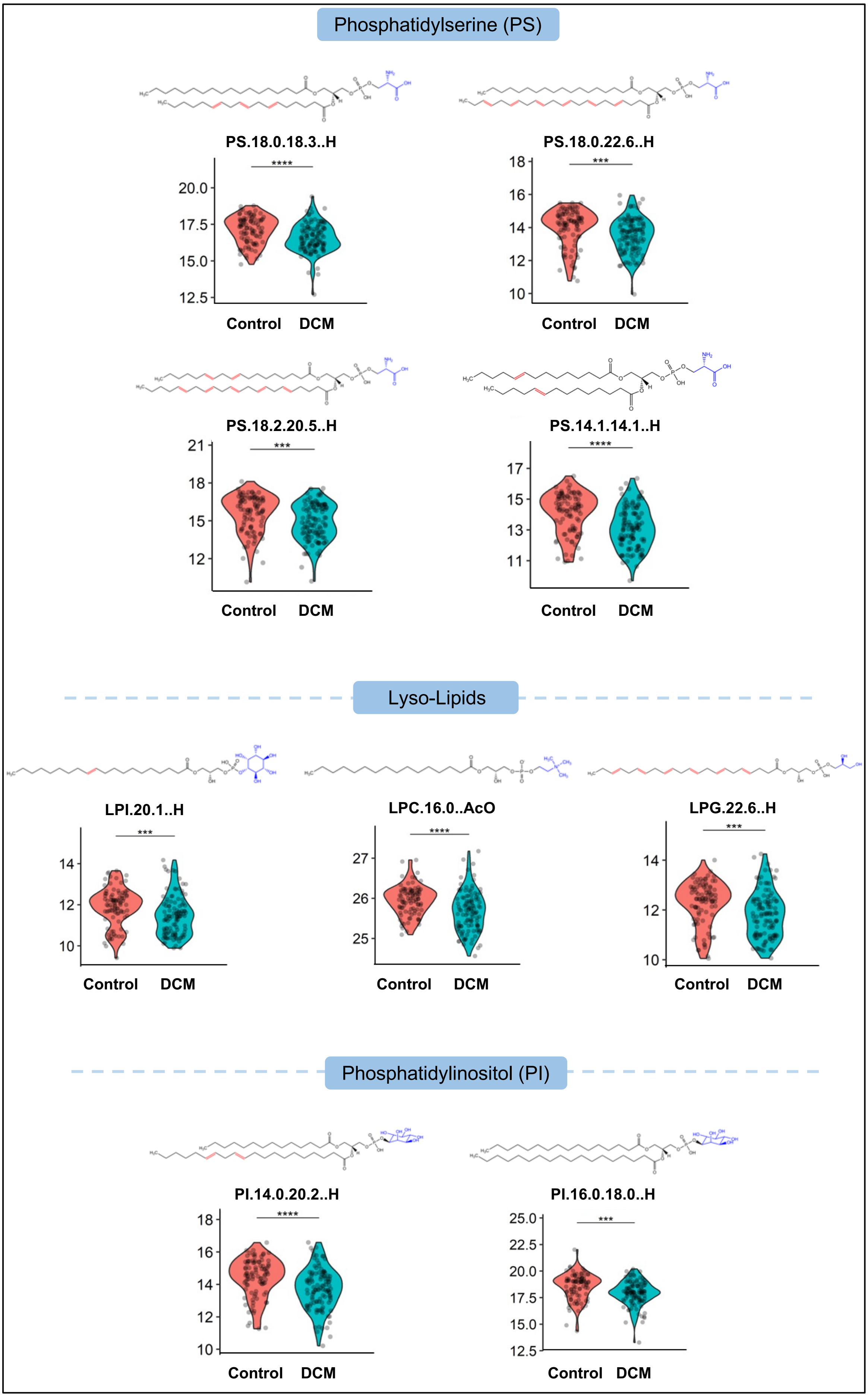

Phosphatidylserine (PS)
PS.18.0.18.3..H
****
20.0
17.5
15.0
12.5
Control
DCM
PS.18.0.22.6..H
***
18
16
14
12
10
Control
DCM
PS.18.2.20.5..H
***
21
18
15
12
Control
DCM
PS.14.1.14.1..H
****
17
15
13
11
Control
DCM
Lyso-Lipids
LPI.20.1..H
***
14
12
10
Control
DCM
LPC.16.0..AcO
****
27
26
25
Control
DCM
LPG.22.6..H
***
14
12
10
Control
DCM
Phosphatidylinositol (PI)
PI.14.0.20.2..H
****
18
16
14
12
10
Control
DCM
PI.16.0.18.0..H
***
25.0
22.5
20.0
17.5
15.0
12.5
Control
DCM

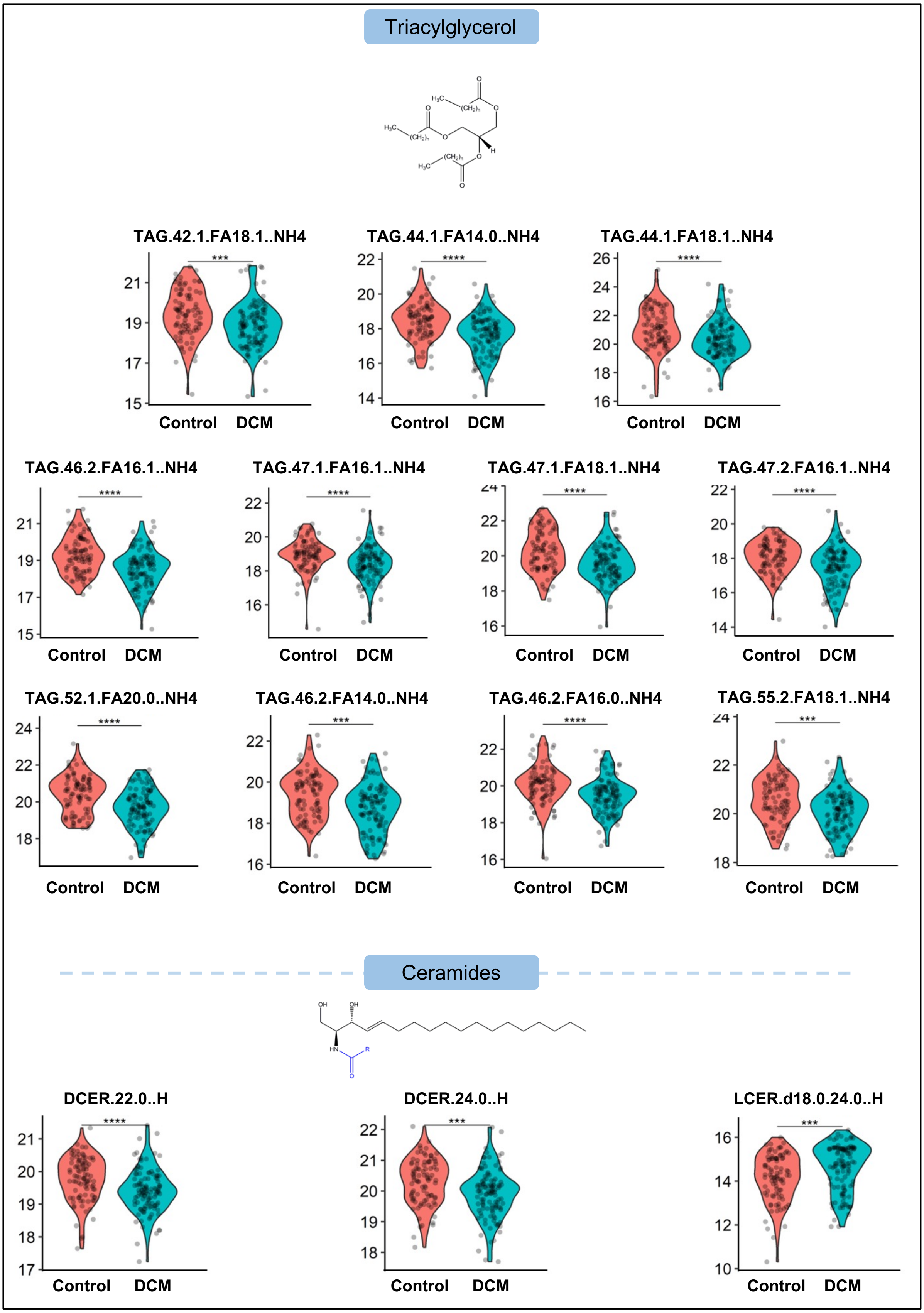


*Supplementary Figure 1: Violin plots of Boruta selected lipids showing significant alteration between DCM and Healthy individuals (*, p<0.05; **, p<0.01; ***, p<0.001; ****, p<0.0001)*

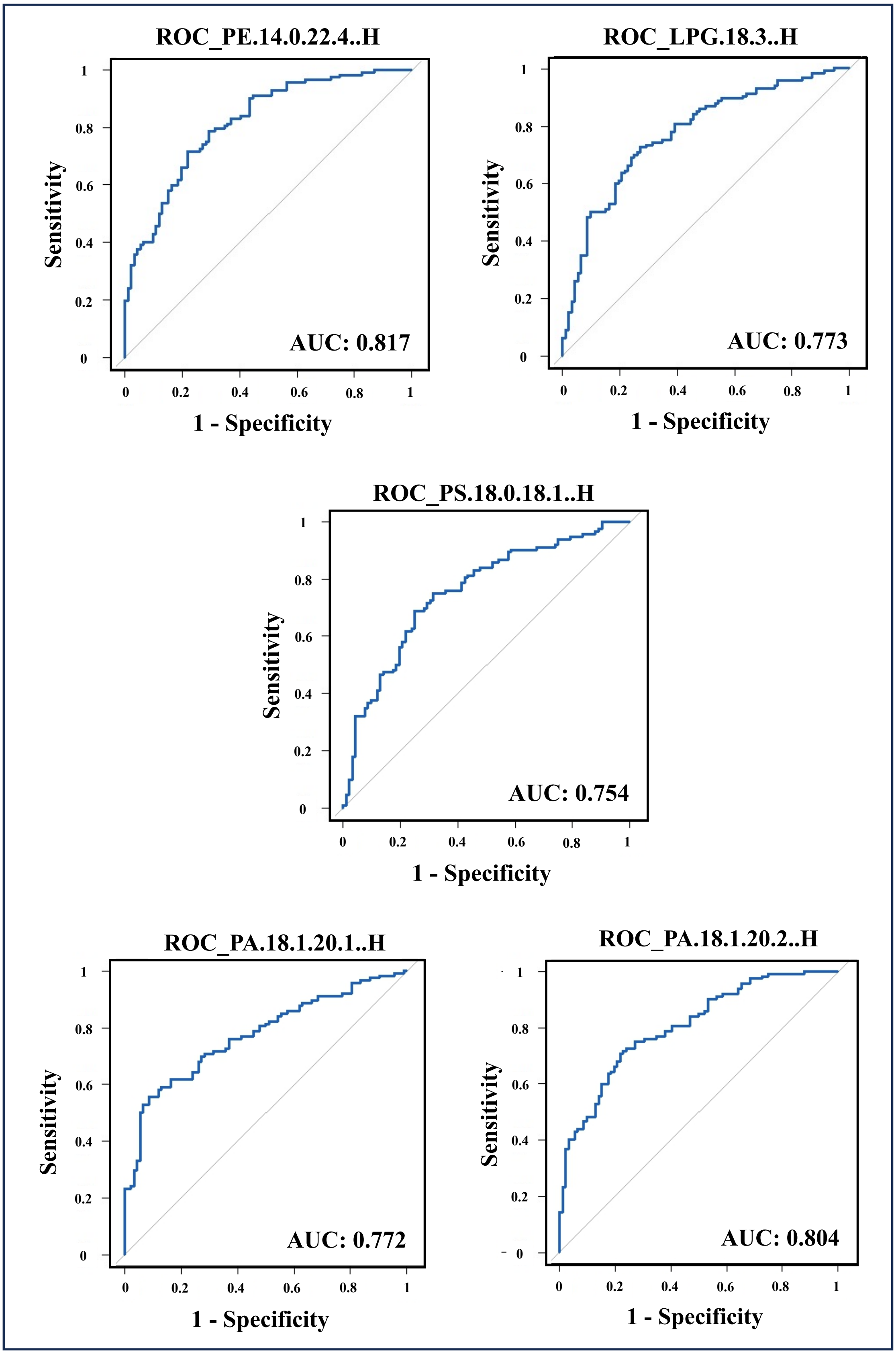
ROC_PE.14.0.22.4..H
AUC: 0.817
Sensitivity
1 - Specificity
ROC_LPG.18.3..H
AUC: 0.773
Sensitivity
1 - Specificity
ROC_PS.18.0.18.1..H
AUC: 0.754
Sensitivity
1 - Specificity
ROC_PA.18.1.20.1..H
AUC: 0.772
Sensitivity
1 - Specificity
ROC_PA.18.1.20.2..H
AUC: 0.804
Sensitivity
1 - Specificity

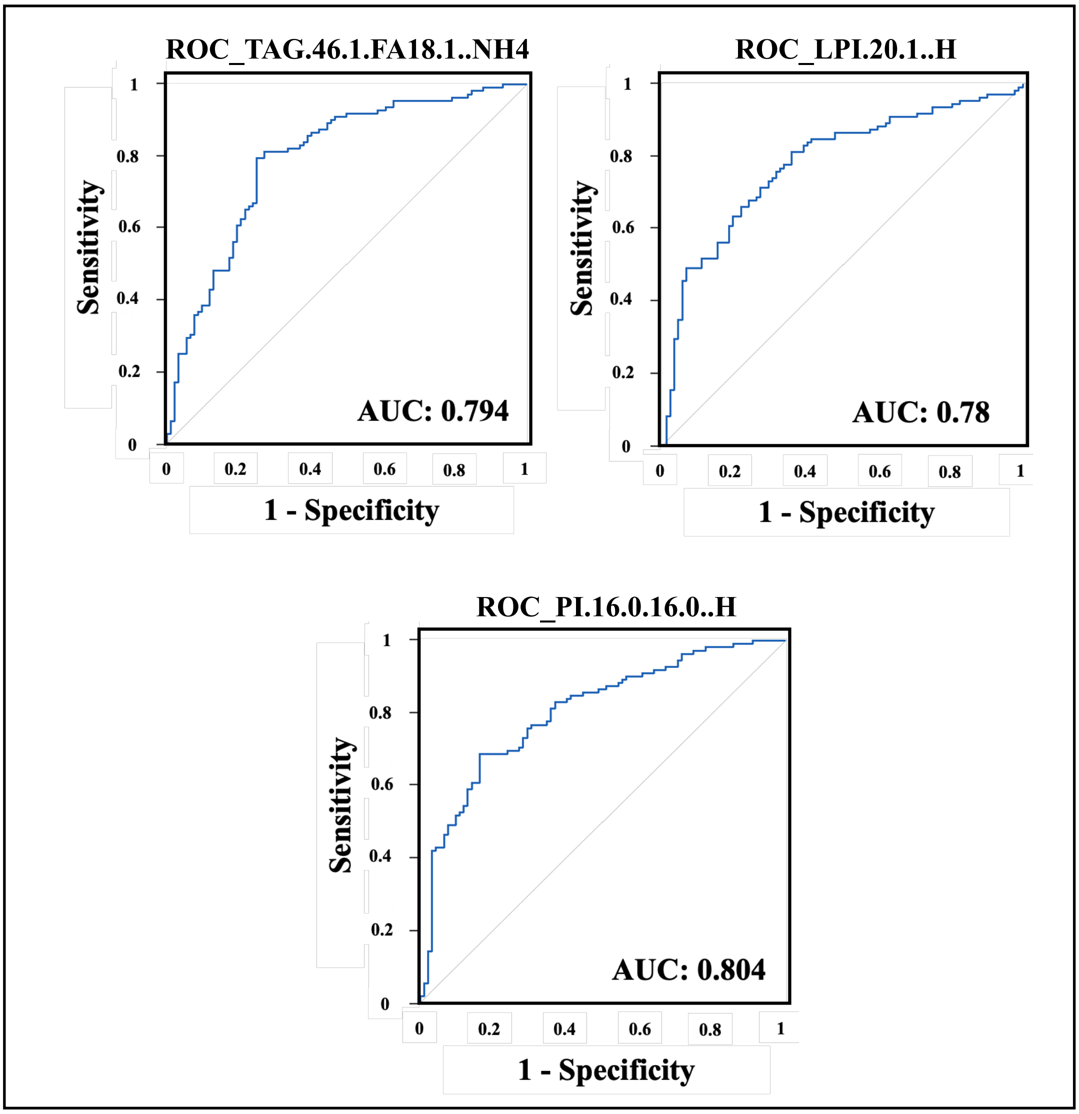


*Supplementary Figure 2: ROC curve analysis of top 5 Boruta selected lipids depicting their predictive accuracy in DCM and Healthy controls*

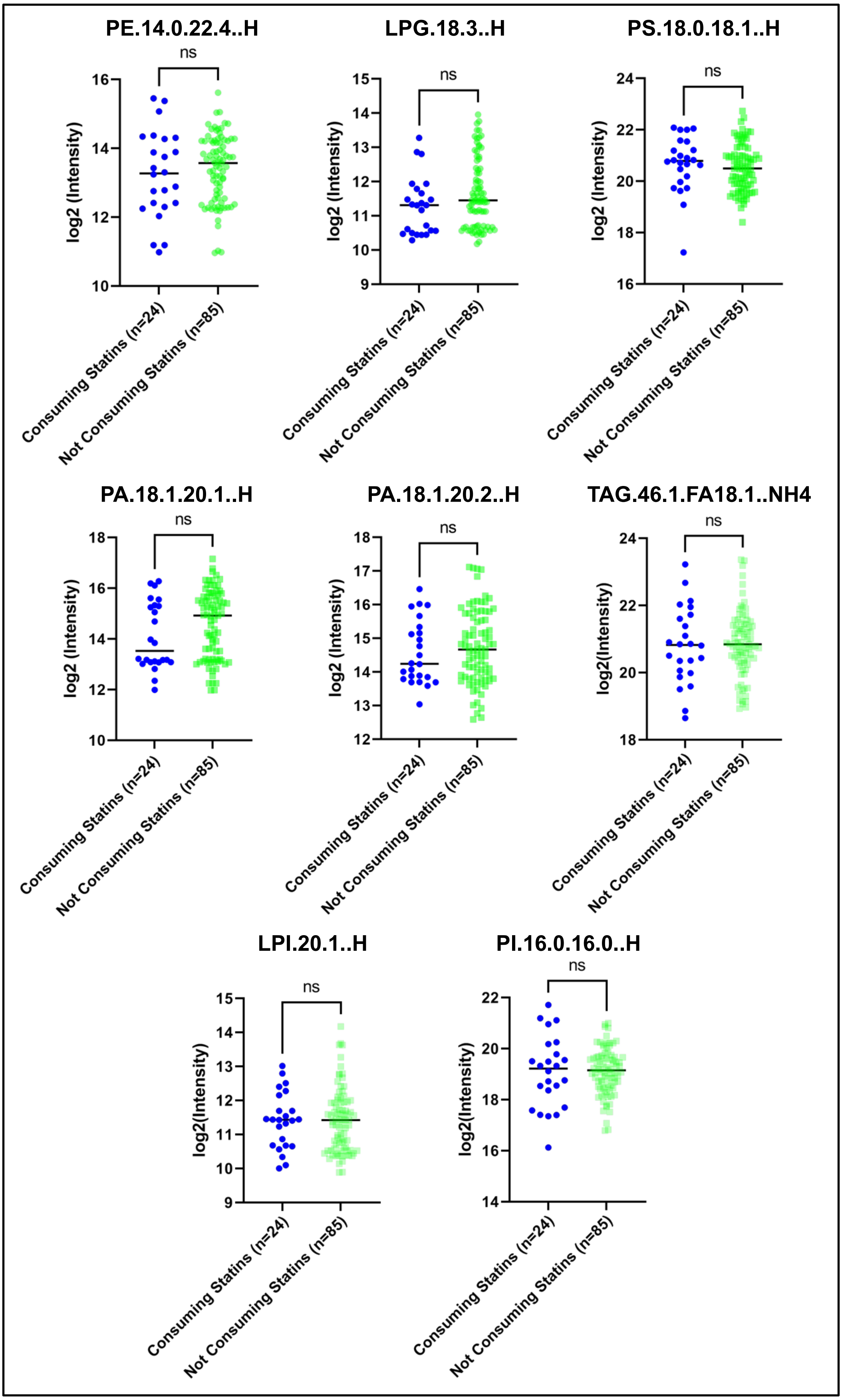


*Supplementary Figure 3: Alteration of top 5 Boruta selected lipid species in DCM patients based on their status of Statin consumption*

*Supplementary Table 2: Boruta-selected significantly altered proteins in DCM patients*

| Uniprot ID\|Gene Name | log fc | -log p | -log $p_{adj}$ | Function |
|---|---|---|---|---|
| P36980\|CFHR2 | 0.285 | 2.538 | 2.538 | Involved in complement regulation. Associates with lipoproteins and may play a role in lipid metabolism[19]. |
| A0A8Q3SIA1\|SUSD1 | 0.279 | 4.821 | 4.520 | Predicted to enable calcium ion binding activity. |
| O14791\|APOL1 | -0.288 | 8.038 | 7.515 | Participates in reverse cholesterol transport from peripheral tissues to the liver[20]. |
| P05452\|CLEC3B | -0.274 | 8.719 | 8.020 | May have some role in myocardial fibrosis[21]. |
| P00738\|HP | 0.386 | 3.903 | 3.824 | Interacts with free plasma Hb to induce recycling of iron and prevents kidney damage. Also functions as antioxidant[22]. |
| P00740\|F9 | -0.281 | 6.886 | 6.488 | Induces intrinsic pathway of blood coagulation by producing the active form of factor X with the help of vitamin K, $Ca^{2+}$, phospholipids and factor VIIIa[23]. |
| P18428\|LBP | 0.274 | 4.112 | 3.957 | Induces inflammatory response[24]. |
| P61769\|B2M | 0.495 | 11.879 | 10.879 | Stimulates exaggerated T-cell response and hyper-activation of RAAS system[25]. |
| P68871\|HBB | -0.509 | 4.341 | 4.119 | Involved in oxygen transportation to various peripheral tissues through blood[26]. |
| P69905\|HBA1 | -0.432 | 3.870 | 3.824 | |

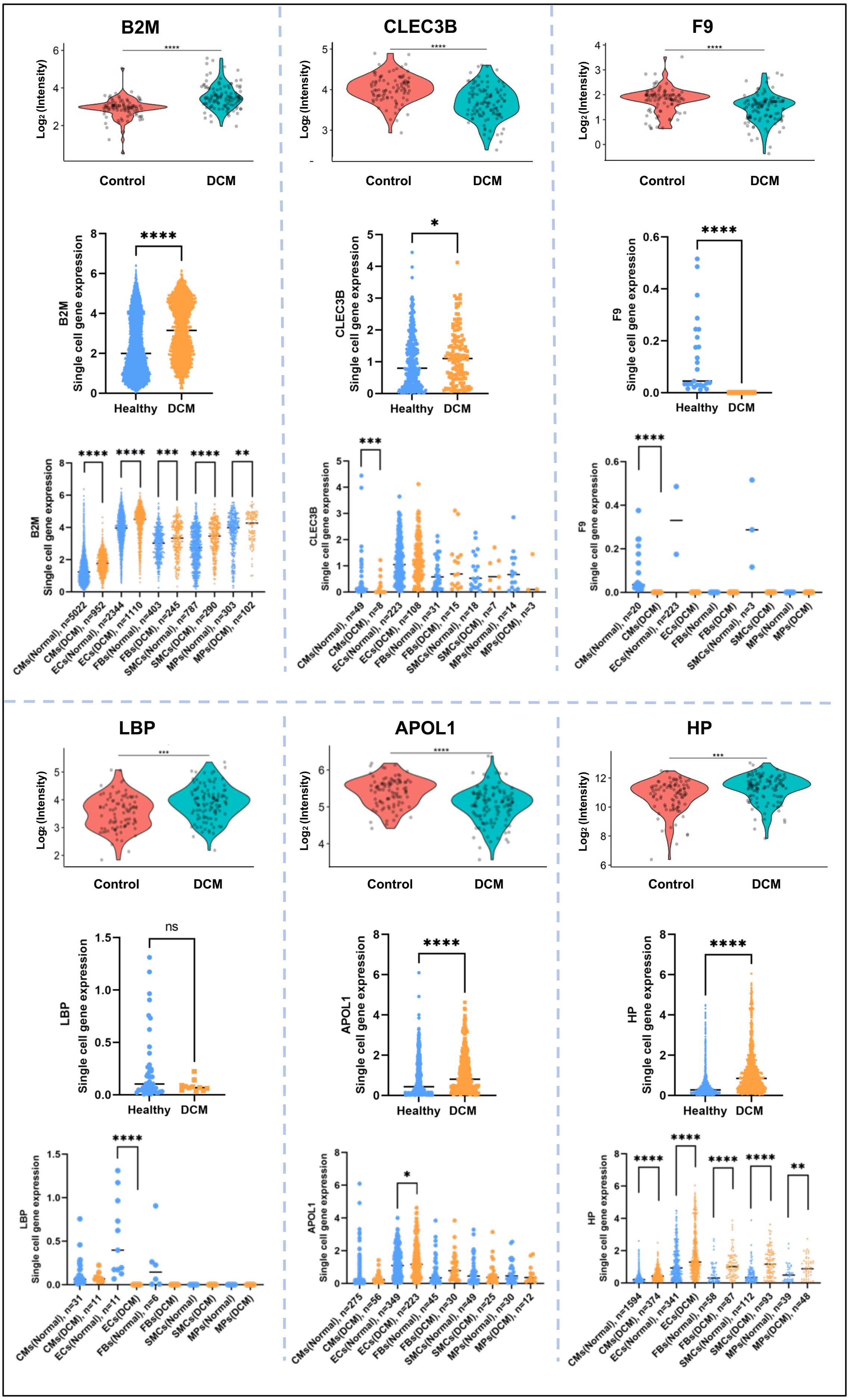
B2M
CLEC3B
F9
LBP
APOL1
HP
Log2 (Intensity)
Control
DCM
Healthy
Single cell gene expression
ns
CMs(Normal), n=5022
CMs(DCM), n=952
ECs(Normal), n=2344
ECs(DCM), n=1110
FBs(Normal), n=403
FBs(DCM), n=245
SMCs(Normal), n=787
SMCs(DCM), n=290
MPs(Normal), n=303
MPs(DCM), n=102
CMs(Normal), n=49
CMs(DCM), n=8
ECs(Normal), n=223
ECs(DCM), n=108
FBs(Normal), n=31
FBs(DCM), n=15
SMCs(Normal), n=18
SMCs(DCM), n=7
MPs(Normal), n=14
MPs(DCM), n=3
CMs(Normal), n=20
CMs(DCM)
ECs(Normal), n=223
ECs(DCM)
FBs(Normal)
FBs(DCM)
SMCs(Normal), n=3
SMCs(DCM)
MPs(Normal)
MPs(DCM)
CMs(Normal), n=31
CMs(DCM), n=11
ECs(Normal), n=11
ECs(DCM)
FBs(Normal), n=6
FBs(DCM)
SMCs(Normal)
SMCs(DCM)
MPs(Normal)
MPs(DCM)
CMs(Normal), n=275
CMs(DCM), n=56
ECs(Normal), n=349
ECs(DCM), n=223
FBs(Normal), n=45
FBs(DCM), n=30
SMCs(Normal), n=49
SMCs(DCM), n=25
MPs(Normal), n=30
MPs(DCM), n=12
CMs(Normal), n=1594
CMs(DCM), n=374
ECs(Normal), n=341
ECs(DCM)
FBs(Normal), n=58
FBs(DCM), n=87
SMCs(Normal), n=112
SMCs(DCM), n=93
MPs(Normal), n=39
MPs(DCM), n=48

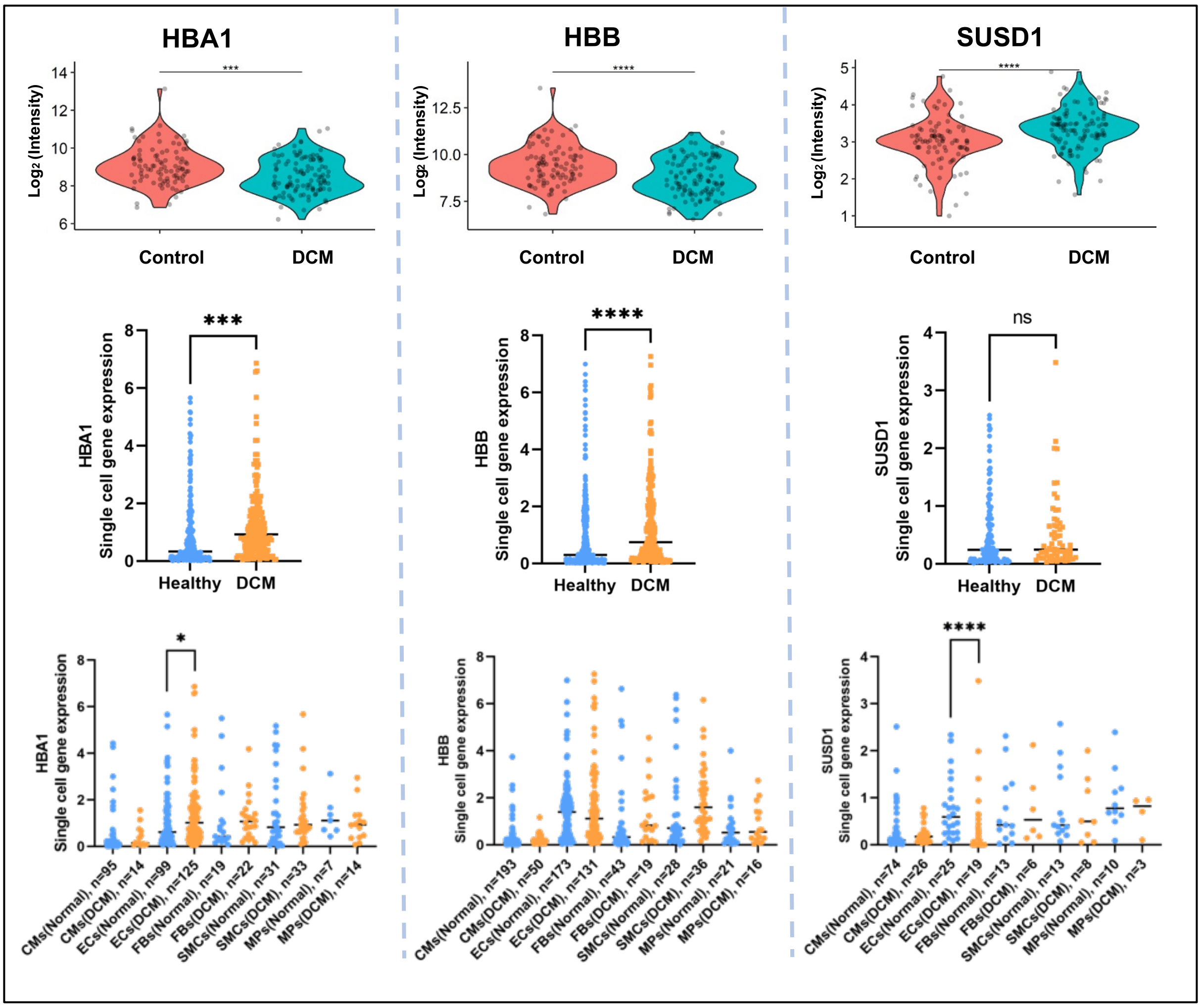


*Supplementary Figure 4: Comparative analysis of the plasma levels of Boruta selected proteins and their tissue expression pattern from single-cell RNA sequencing data (*, $p<0.05$; **, $p<0.01$; ***, $p<0.001$; ****, $p<0.0001$)(B2M=beta 2 microglobulin; HP=Haptoglobin; F9=Coagulation factor IX; LBP=Lipopolysaccharide binding protein; APOL1=Apolipoprotein L1; CLEC3B=Tetranectin; HBA1=Hemoglobin subunit alpha; HBB=Hemoglobin subunit beta; SUSD1=Sushi domain nidogen containing protein; CM=Cardiomyocytes; EC=Endothelial Cells; FB=Fibroblast; SMC=Smooth muscle cell; MP=Macrophages)*

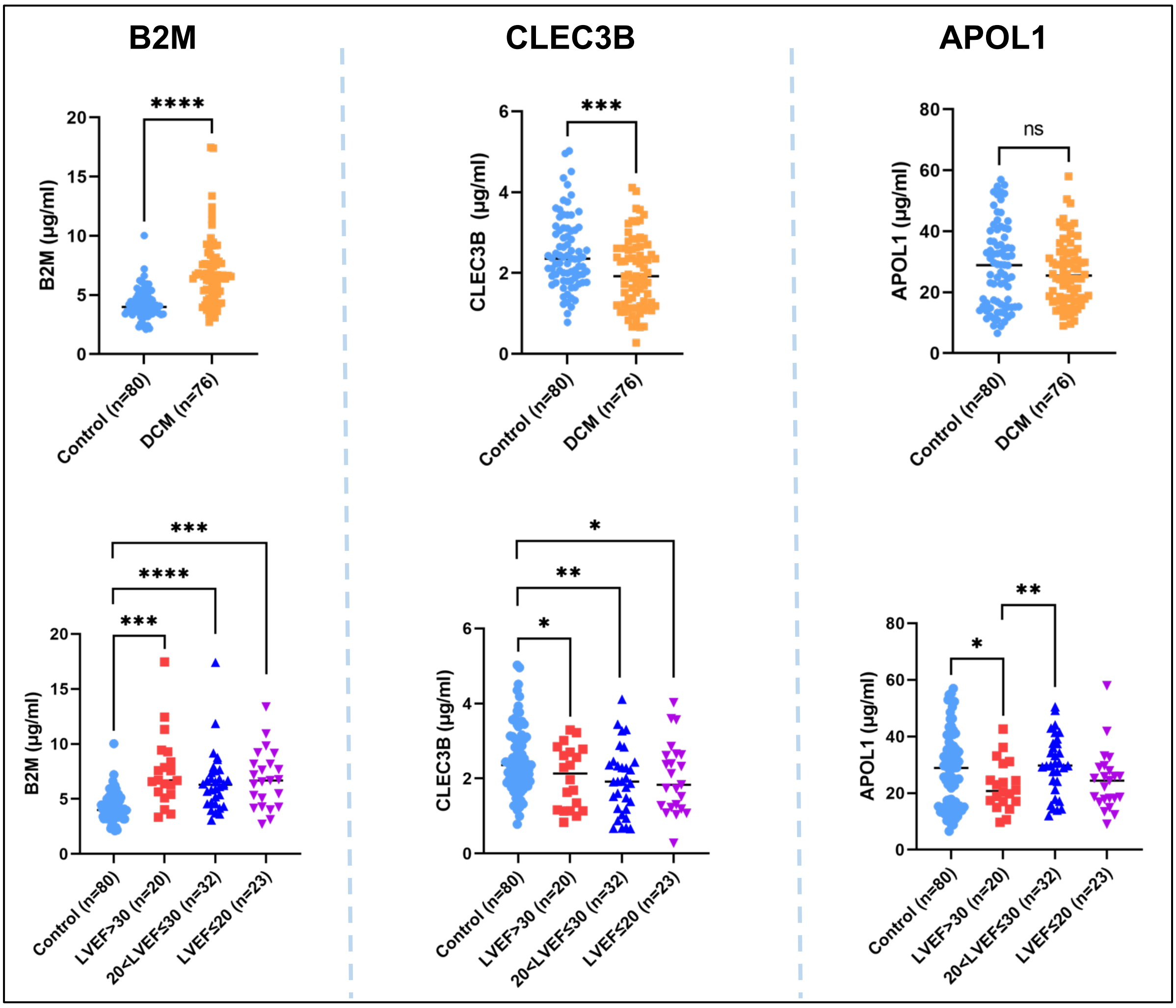


*Supplementary Figure 5: Dot-plot representation of ELISA data of 3 Boruta selected proteins and comparative analysis of the plasma levels of those proteins across different LVEF quartiles. ns=Non-significant, *, p<0.05, **, p<0.01, ***, p<0.001, ****, p<0.0001.*

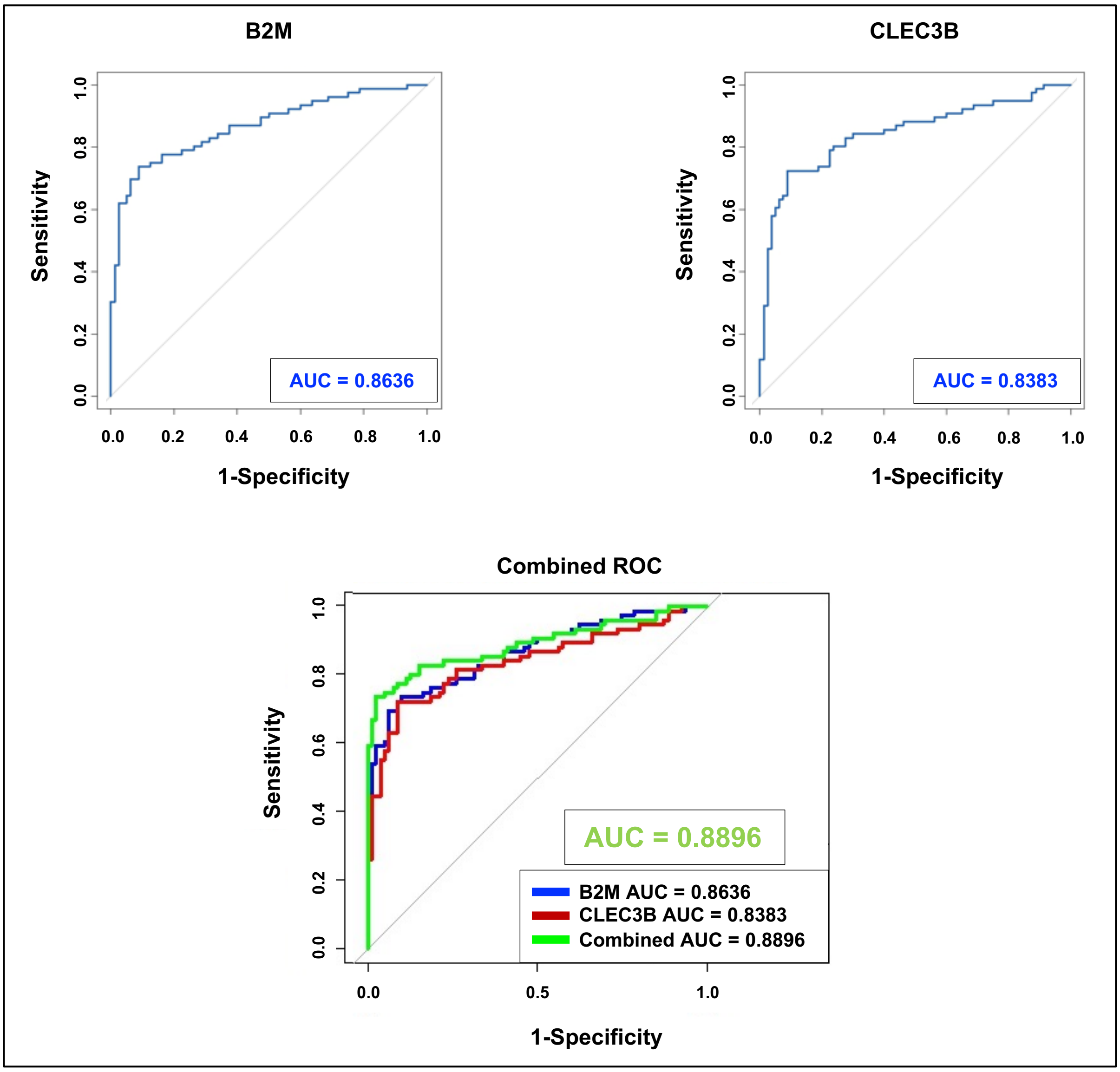


*Supplementary Figure 6: ROC curve analysis of 2 validated proteins (B2M and CLEC3B) through ELISA depicting their predictive accuracy in DCM and Healthy controls*
